# Exploring “dark matter” protein folds using deep learning

**DOI:** 10.1101/2023.08.30.555621

**Authors:** Zander Harteveld, Alexandra Van Hall-Beauvais, Irina Morozova, Joshua Southern, Casper Goverde, Sandrine Georgeon, Stéphane Rosset, Michëal Defferrard, Andreas Loukas, Pierre Vandergheynst, Michael M. Bronstein, Bruno E. Correia

## Abstract

*De novo* protein design aims to explore uncharted sequence-and structure areas to generate novel proteins that have not been sampled by evolution. One of the main challenges in *de novo* design involves crafting “designable” structural templates that can guide the sequence search towards adopting the target structures. Here, we present an approach to learn patterns of protein structure based on a convolutional variational autoencoder, dubbed Genesis. We coupled Genesis with trRosetta to design sequences for a set of protein folds and found that Genesis is capable of reconstructing native-like distance-and angle distributions for five native folds and three novel, so-called “dark-matter” folds as a demonstration of generalizability. We used a high-throughput assay to characterize protease resistance of the designs, obtaining encouraging success rates for folded proteins and further biochemically characterized folded designs. The Genesis framework enables the exploration of the protein sequence and fold space within minutes and is not bound to specific protein topologies. Our approach addresses the backbone designability problem, showing that structural patterns in proteins can be efficiently learned by small neural networks and could ultimately contribute to the *de novo* design of proteins with new functions.

## Introduction

Evolution is a gradual and slow process that has only sampled a *minuscule* fraction of the possible protein sequence space^1^. In many instances, natural sequences collapse into three-dimensional (3D) structures that can be categorized into a finite set of protein folds. To explore novel sequences that fold into well-defined 3D-conformations outside the natural repertoire and likely amenable to new functionalities, *de novo* protein design strategies have been developed^2^. Most structure-based *de novo* protein design approaches rely on a two-step process where (1) the protein fold is outlined and corresponding backbones are generated, and (2) amino acid (AA) sequences are searched for to fit the generated backbones. Machine learning (ML) techniques have been explored to assist both steps.

Despite recent successes^3–7^, robust *de novo* design remains challenging. One of the long-standing challenges in *de novo* design is the backbone generation stage where physically unrealistic conformations lead to sequences that fail to fold experimentally. Designable backbones present optimal secondary structure configurations that pack into tertiary conformations such that they are effectively realizable with the 20 natural AAs^8–12^. It has been observed that the number and variety of energetically favorable sequences that fit onto a certain protein structure is linked to the designability of the backbone^8,12^. However, quantitatively describing designability remains a largely unsolved problem because it includes properties that are difficult to measure, such as fold specificity^9,13^, or native-like structural arrangements^14^, and therefore it is challenging to optimize for such a property.

Physics-based *de novo* design methods attempt to capture backbone designability by explicitly formulating empirical rules based on loop lengths and structure that embed the packing of local tertiary motifs to secondary structure elements (SSEs) and result in strongly linked sequence-structure relationships^15,16^. These rules, together with fragment assembly protocols^17^, led to the design of “ideal” protein folds with small loops and regular SSEs.

Recent advances in deep neural networks (DNNs) combined with the availability of large-scale protein sequence and structural datasets have enabled highly accurate structure prediction from sequence^18,19^. Strikingly, structure prediction networks can be “reversed” for the protein design task. For example the transform-restrained Rosetta (trRosetta) or AlphaFold (AF) networks can be employed for fixed backbone design via backpropagating gradients from the target structure to the sequence. Employing structure prediction methods in reverse has the effect of implicitly optimizing over the full sequence and structure landscape^20,21^ i.e., searching for the lowest-energy sequence while maximizing the probability of the target structure relative to all other conformations, which improves fold specificity.

Other methods use structure-based context given by the protein backbone to generate sequences. Popular approaches utilize message-passing or 3D convolutional neural networks to encode structural features of protein backbones, and subsequently decode optimal AA sequences for the given input backbone conformation^22–24^. Encouragingly, DNNs are able to design new sequences for a target structure within minutes on modern graphical processing units (GPUs), enabling diverse and high-throughput sampling of the design space^25^.

DNNs can also be applied to the *de novo* protein design task where novel proteins are created from scratch. One approach involves trRosetta where novel proteins are hallucinated by using a specific loss that maximizes the contrast between random (background) and native distance distributions^26^. Recently, DNNs based on denoising diffusion probabilistic models, such as Chroma^27^ and RFdiffusion^28^, have demonstrated the ability to generate novel proteins starting from Gaussian noise. Remarkably, the two latter methods can be guided to adhere to specific input constraints, allowing for the generation of small to large protein domains with desired shapes, both symmetric and asymmetric assemblies, and various other protein characteristics.

While powerful in generating medium to large novel protein folds, many DNNs encounter challenges in generating small new protein domains not observed in nature i.e., domains with novel configurations or wiring of the SSEs^26,28^. For example, trRosetta-based hallucination often produces protein backbones with a TM-score exceeding 0.6 to the Protein Data Bank (PDB^29^), indicating that it mainly recapitulates pre-existing folds^26^. These observations imply that computationally generating small novel protein domains that ultimately work experimentally continues to present a challenging endeavor.

Here, we investigate current challenges in designing small novel protein folds and propose a new approach based on learning 3D structural patterns from protein backbones, referred to as backbone designability^8,10,11^. Our proposed workflow involves the representation of protein folds in a string format that is projected into 3D-space as a protein “sketch” (Fig. 1A and Sup. Fig. 1). A sketch is constructed using a low-resolution placement of SSEs to approximate and facilitate the fold search. To improve the designability of the sketch, we use a convolutional variational autoencoder (VAE), dubbed *Genesis*, to encode the distances and orientations of sketches into a compact latent representation that can be sampled and subsequently decoded into native-like distance and orientation probabilities (Sup. Fig. 1A). The refined distance and orientation probabilities function as a target template to guide trRosetta in generating new backbones and sequences that adhere to the template restraints (Fig. 1A, Sup. Fig. 2 and Sup. Fig. 3).

**Figure 1.**
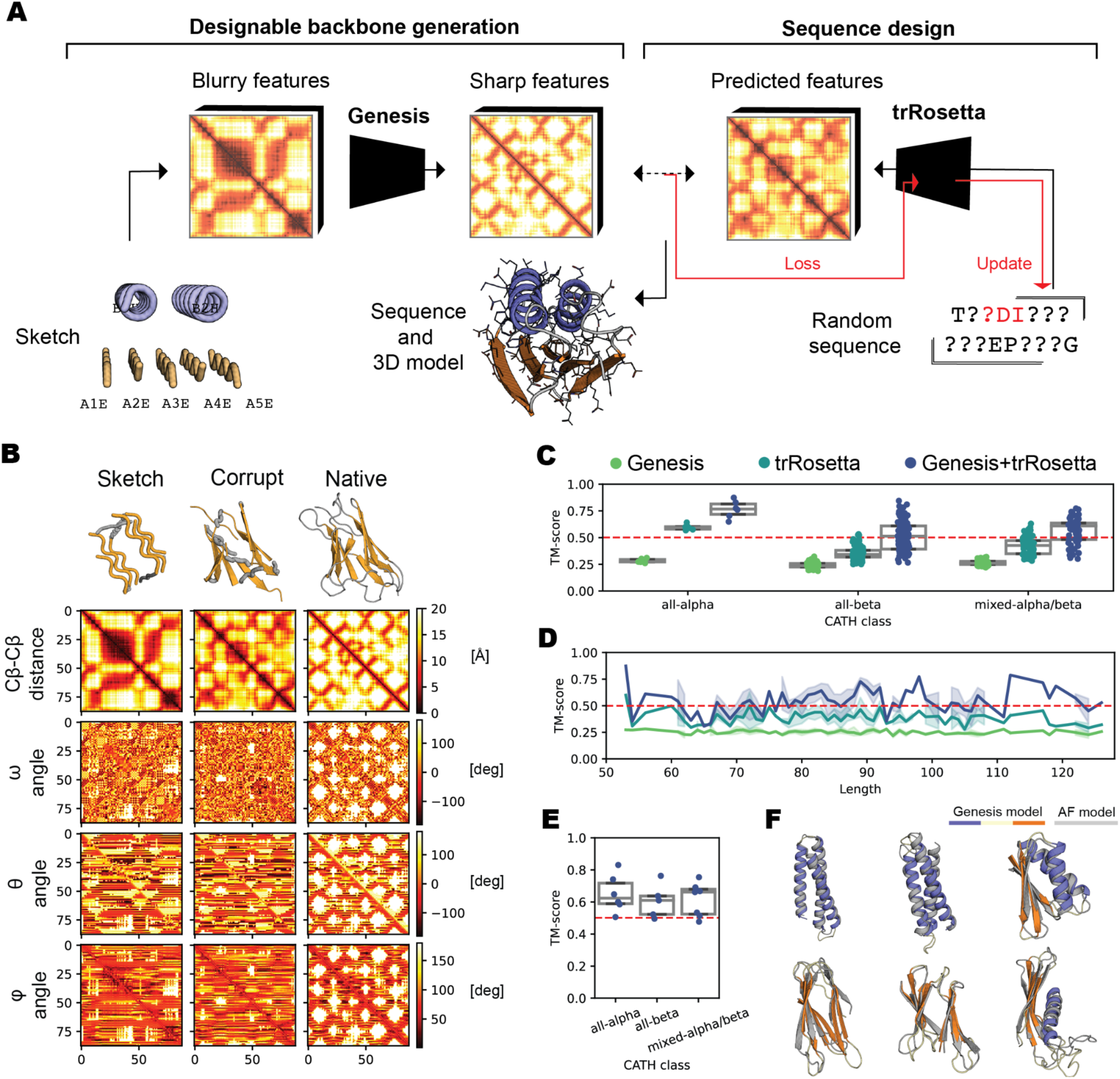
Overview and benchmarking of the Genesis-trRosetta *de novo* protein design pipeline. **A:** A protein sketch, which defines the SSE content and the fold, is specified as input to Genesis. The sketch displays blurry structural features (distances and orientations) that are denoised, sharpened and complemented by Genesis. The trRosetta fixed backbone design strategy is used to design sequences obeying the refined structural features yielding sequences and 3D protein models with the target specified shape. **B:** Comparison of the structure feature maps between a sketch, corrupted structure, and native structure. The Sketch displays coarse-grained features of the protein structure, while the native structure has fine-grained features with many structural details. **C:** Ablation experiments on randomly selected training set examples shows that Genesis and trRosetta modules are required for the best performance. **D:** The Genesis-trRosetta performance is sequence length independent, with a maximum length of 128 (determined by the size of the input into the model). **E:** Fold recovery experiment of 18 folds with SCOPe families and input sketches not included in the training set shows the generalizability of Genesis-trRosetta. **F:** AF predictions and structural comparisons of test examples.

We performed a comprehensive battery of computational tests to further understand the strengths and weaknesses of our computational pipeline. Furthermore, we sought to experimentally evaluate the capabilities of our design approach by computationally generating sequences for five native folds and three “darkfolds” with different secondary structure and loop lengths and assessing their folding using protease resistance assays for 550 designs and biochemical characterization analysis of 39 individual designs.

### A computational pipeline for enhanced backbone designability

We developed a *de novo* design workflow based on deep learning modules where the Genesis module (Fig. 1A) explicitly learns and incorporates native-like structural features into low resolution sketches of protein structures. Genesis itself relies on a convolutional variational autoencoder (VAE) architecture that is trained to transform a large dataset of small sketches into their respective native structures (Sup. Fig. 1A). The developed convolutional VAE operates on pairwise distances and orientations rather than atomic coordinates (see Methods for implementation details and data encoding). Importantly, distances and orientations are invariant with respect to translation and rotation which ensures stable performance in the presence of transformations of the data input under the special Euclidean group SE(3)^30^. Our VAE is conditioned on real-valued distances and orientations derived from the sketches, and predicts distance-and orientation probabilities of native-like conformations from the latent conditional distribution. We integrated Genesis with the trRosetta fixed backbone design method, which performs the sequence search guided by the distance and orientation probabilities. It is worth mentioning that other tools that operate on distances and orientations such as AF or RosettaFold could potentially also be adapted and utilized for the sequence design.

To test the generalization capabilities of Genesis, we split our data into a training and a test set based on the Structural Classification of Proteins — extended (SCOPe) definition (see Methods for details and Sup. Fig. 4). The sketches display significant differences in both pairwise distance-and orientation maps when compared to those of native protein structures as depicted in Fig. 1B, and hence converting a sketch into a native-like backbone is challenging. This can be attributed to the fact that in a sketch, the SSEs are (1) naively layered without considering any physical constraints, and (2) loops are absent and modeled as random residues positioned in a straight line connecting two SSEs.

We randomly selected 273 diverse all-alpha, all-beta, and mixed-alpha/beta folds from the training set and conducted ablation studies in the different modules of the pipeline, shown in Fig. 1C (details in Methods). We compared the Genesis-only, trRosetta-only and Genesis-trRosetta frameworks using a self-consistency TM-score e.g., aligning the structural predictions of the designed sequences onto the native target fold.

The Genesis-trRosetta protocol improves structural fold recovery for each SCOPe class and is capable of generating multiple backbones that have median TM-scores above 0.5 (Fig. 1C). For all-alpha topologies the ablated frameworks (Genesis-only and trRosetta-only) reach median TM-scores around 0.25 and 0.6 respectively, whereas the full Genesis-trRosetta protocol reaches a median TM-score of 0.75 (Fig. 1C). For all-beta and mixed-alpha/beta folds, both ablated frameworks have median TM-scores below 0.5 and only a few sampled backbones have TM-scores around the 0.5 threshold (Fig. 1C). Using the complete Genesis-trRosetta framework roughly 40-50% of the sampled backbones have TM-scores above 0.5, with the best recoveries reaching TM-scores around 0.75 (Fig. 1C).

We further investigated if there is a sequence length dependence for proteins between 50 and 128 residues (Fig. 1D). We observe that the ablated protocols perform poorly compared to the full Genesis-trRosetta protocol and that folds with 110-120 residues seem to have higher recovery rates (Fig. 1D). Many folds with poor TM-score are examples that are difficult to define as a sketch e.g., fully beta proteins such as right-handed beta-helix folds, alpha/beta propellers, or beta-Prisms which consist of three beta-sheets roughly packing into prism-resembling fold.

We next tested if the Genesis-trRosetta framework is capable of recovering folds that were not included in the training set (Fig. 1E, F). The Genesis-trRosetta framework was used to sample backbones for 18 test structures with sketches plus SCOPe families that were not included in the training set (details in Methods). The results indicated that Genesis-trRosetta is capable of generating backbones with TM-scores between ∼0.5-0.8 for all-alpha, all-beta and mixed-alpha/beta structures, indicating that the framework can generalize outside the training set (Fig. 1E, F).

In summary, these results provide evidence that the Genesis-trRosetta pipeline enhances the backbone designability of low resolution protein sketches generating sequences that are predicted to adopt the target fold. In addition the pipeline has also shown good generalizability towards protein shapes that are absent from the training set.

### Large-scale *de novo* design of native topologies

To showcase the Genesis-trRosetta *de novo* design framework, we set out to design five different topologies. We sampled a two-layer mixed alpha/beta Ubiquitin-like fold, where four strands are packed against a helix, and a three-layer mixed alpha/beta Rossmann fold with a central four stranded beta-sheet and two exterior packing helices on both sides. We additionally challenged the framework by designing two different two-layer beta-sandwiches, an Immunoglobulin (Ig) -like fold and a Jellyroll fold. Finally, we designed sequences that adopt the Top7 fold^31^, a novel fold that is not present in the natural repertoire and can represent a generalization test to our method. As an indication of fold designability, we used the relative contact order^32^ for each of the above-mentioned folds to measure the locality of the structural contacts. Contact order is defined as the average sequence separation between residues that are in contact within the folded structure of the protein. Importantly, proteins with high contact order tend to fold slower than those with low contact order^32^, and remain challenging to design. Ubiquitin-like, Rossmann and Top7 folds have a contact order of around 0.15 indicating a larger presence of local contacts. On the other hand, Ig-like and Jellyroll folds exhibit greater non-locality of structural contacts having relative contact orders of approximately 0.20 and 0.26, respectively.

For each of the topologies, we did not have prior knowledge about secondary structure-and loop length distributions that could successfully be assembled into designable backbones. Therefore, we adopted an “exploration-exploitation” strategy and performed a two-stage sampling approach (Sup. Fig. 5 and Sup. Fig. 6): (1) We sampled over 20-30 secondary structure and loop length combinations which we refer to as “candidate search stage” to generate candidate templates with potentially designable length combinations (Sup. Fig. A); (2) candidate templates that fulfilled the self-consistency structure prediction metrics (Sup. Fig. 7) were carried to a “production stage” where around 20,000 sequences and corresponding 3D models were generated (Fig 2A, Sup. Fig. 8 and Methods for detailed explanation of both stages).

**Figure 2.**
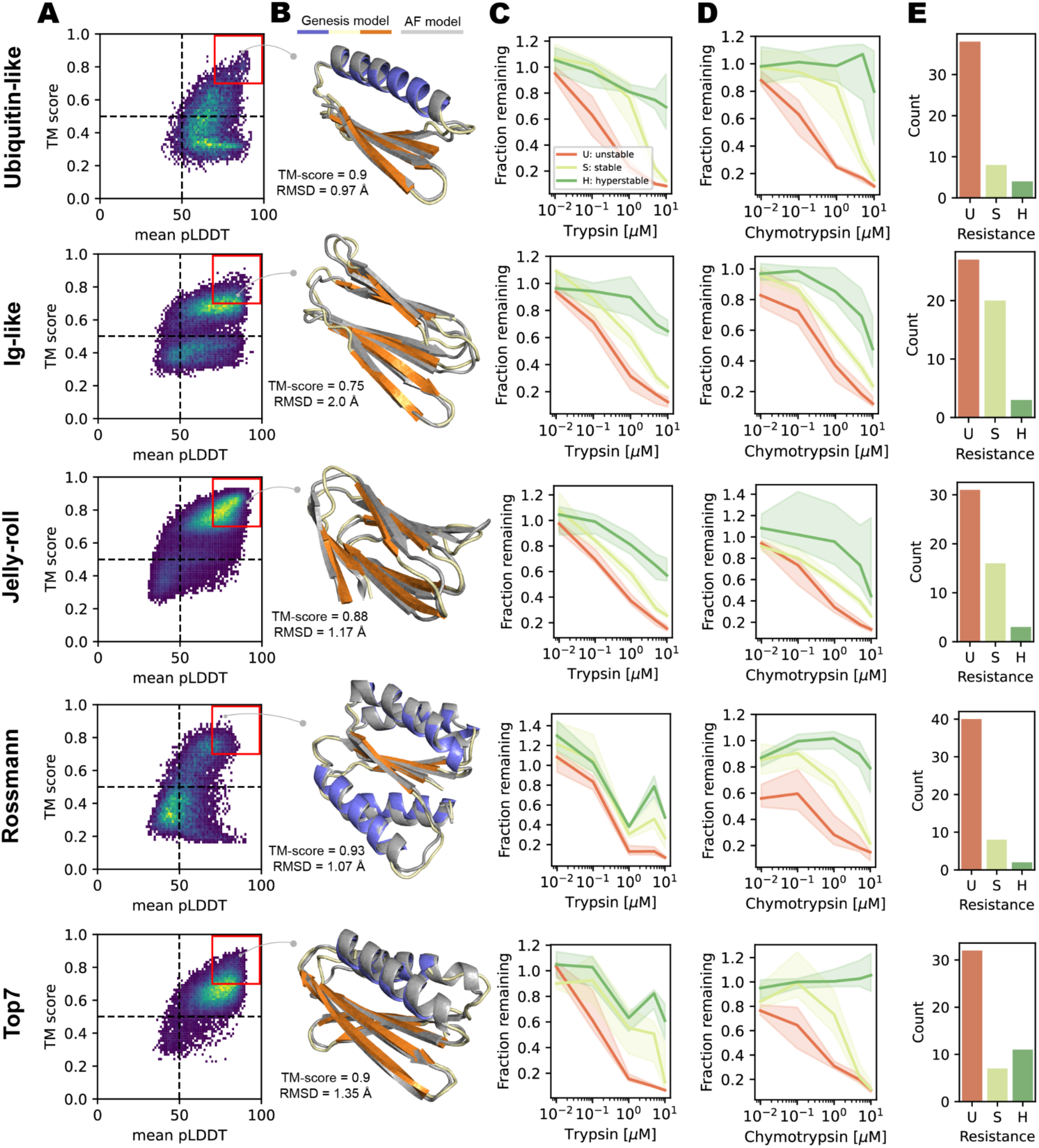
Computational selection and high-throughput experimental protease resistance screening of designs with native folds. **A:** *In silico* design success was determined by AF structure predictions. Designs with both a pLDDT (> 70) and high TM-scores between the Genesis and the AF model (> 0.7). **B:** Genesis and AF structural model examples for each fold. **C, D:** Results of an experimental high-throughput enzymatic resistance assay on yeast (using Trypsin and Chymotrypsin proteases). The designs were subjected to increasing concentrations of Trypsin and Chymotrypsin proteases, leading to preferential degradation of unfolded designs. For each design, protease resistance EC_50_s were calculated and the designs categorized as high resistance (H, protease resistance EC_50_ > 5 μM), medium resistant (M, 5 μM > protease resistance EC_50_ > 1 μM), and low resistance (L, protease resistance EC_50_ < 1 μM). The median curve for each stability category is graphed with a confidence interval (CI) of 95% (shaded regions). **E:** Overall stability of the designs for Trypsin + Chymotrypsin, where designs were required to show either high or medium resistance in both protease digestions.

To evaluate the performance of our *de novo* design pipeline, a subset of 250 designs (50 per fold) were selected based on the self-consistency TM-score between the model and the AF prediction and the AF pLDDT score (Fig. 2A, B and Sup. Fig. 8). These designs were screened on the surface of yeast to assess their folding using a proteolytic digestion assay as a proxy (Sup. Fig. 9). The yeast library was treated with varying concentrations of protease and analyzed by deep sequencing to identify folded and unfolded designs (for more details see the Methods section).

A folding score (digestion resistance) for trypsin and chymotrypsin was calculated for each design providing an estimate of the folding of the design through the amount of enzyme needed to digest half of the population monitored at the yeast surface^33^ (details in Methods section). Any calculated score above 10 μM is shown as >10 μM, given the upper limit of our assay. There was a wide range of protease resistances measured in our assay. The designs were categorized into three groups according to protease resistance (high resistance (`H`), medium resistance (`M`) and low resistance (`L`)) for analysis (Fig. 2C, D). Each fold type had at least one highly digestion resistant design, with some folds having more designs with high and medium digestion resistance than others (Fig. 2C, D). We additionally tested the stability of native protein sequences for the designed folds (all but one, see Methods). We found that native sequences were within high to medium digestion resistance in our assay, indicating that our designs were on par with native protein sequences (Sup. Fig. 10).

The overall trends (Fig. 2E) of this high-throughput assessment revealed that in each fold a small percentage were highly resistant (∼5-10% of the designs), with the Top7 fold showing the highest percentage of highly folded designs suggesting that such fold may be amenable to host a large number of possible sequences without compromising its stability. Two native beta-sandwich folds that have been particularly difficult for computational design approaches, which has been hypothesized to occur due to the non-local contacts within the folds^34,35^, were selected. According to our folding assessment such folds did not show any reduced success rates having many designs in the intermediate folding category, 20% and 15% of the designs for the Ig-like fold and the Jellyroll, respectively (Fig. 2E). Overall, these results show that our design approach was successful even for folds that are typically considered of high difficulty.

Next, we sought to biochemically characterize several of the medium and highly digestion resistant designs. We obtained synthetic genes for 44 designs, which were expressed and purified from *E. coli* (see Methods for more details). Characterization data for one design of each fold type is shown in Fig. 3.

**Figure 3.**
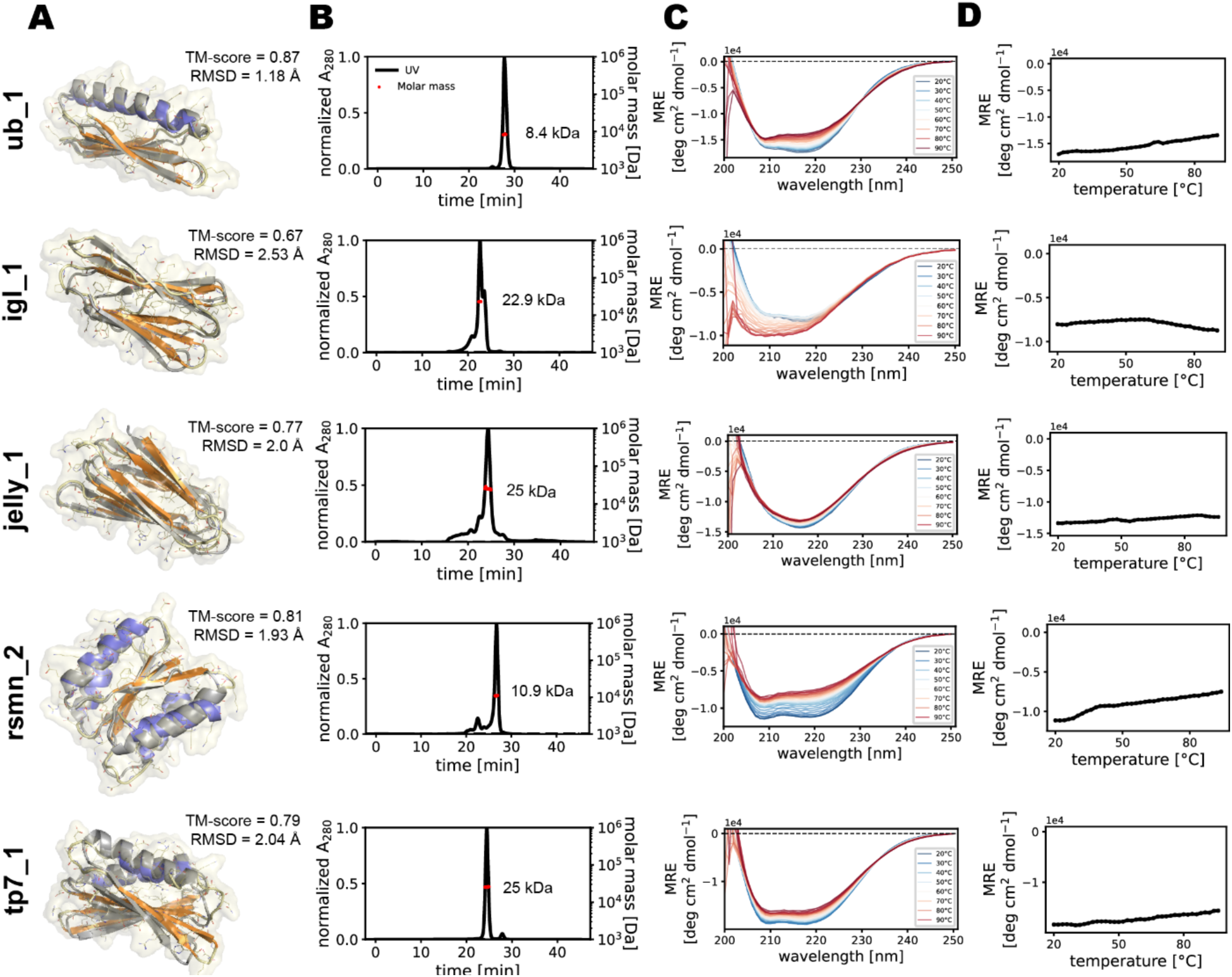
Experimental characterization of the Genesis-trRosetta designs for native folds. **A:** Structural models of the designs for each native fold. **B:** SEC-MALS elution profiles showing the oligomeric state of the designs in solution. Red dots show the molecular weight as determined by SEC-MALS. The theoretical molecular weights for a monomer of the designs are: ub_1: ∼7.5 kDa, igl_1: ∼8.4 kDa, jelly_1: ∼10.4 kDa, rsmn_2: ∼12.5 kDa, tp7_1: ∼11.4 kDa. **C:** Circular dichroism spectroscopy at two different temperatures showing that the designs adopt folded structures in solution. **D:** Thermal denaturation experiments using circular dichroism spectroscopy to determine melting temperatures (T_m_) of the designs.

For the Ubiquitin-like fold, we tested 5 designs out of which 3 were monodisperse in solution and folded according to CD spectroscopy (Sup. Fig. 11). Two of the designs showed the expected monomeric species in solution (ubi_1 and ubi_2) and one was a dimer (ubi_3) (Fig. 3 and Sup. Fig. 11B). Comparisons between the Genesis-trRosetta models with AF were in close agreement showing RMSDs below 1.5 Å (Sup. Fig. 11A).

For the Ig-like fold we attempted to express 11 designs and we obtained two that were purifiable and folded (Sup. Fig. 12). igl_1 shows two oligomeric species-and igl_2 shows a dimer in solution (Fig. 3 and Sup. Fig. 12B). Both designs have a CD signature similar to those of native Ig-like folds and were thermostable (Sup. Fig. 12C, D). The RMSDs of the Genesis-trRosetta models with AF ranged between 1.8-2.5 Å showing a very good agreement between both models (Sup. Fig. 12A).

We next tried to design the Jellyroll fold, one with non-local SSEs connectivities. We expressed 10 designs and only jelly_1 was monodisperse and folded in solution (Fig. 3). The design was a dimer in solution and presented a CD spectrum with a beta-sheet signature as well as high thermal stability (Fig. 3B-D). The designs of this fold were the least successful of the whole set attempted, highlighting the difficulties of topologies with non local contacts.

Next, we tested our design pipeline on a Rossmann fold. We obtained 3 out of 9 designs that were folded in solution (Sup. Fig. 13), from which 2 were monomeric and the remaining was dimeric (Fig. 3 and Sup. Fig. 13B). The AF predictions are in agreement with Genesis-trRosetta pipeline models showing RMSDs within 1.9 to 2.6 Å (Sup. Fig. 13A).

Lastly, we attempted to design a topology that has not been observed in nature, Top7. For this fold, 3 out 9 designs were soluble, folded and with all of them showing high thermostability and two being dimers in solution while the remaining is monomeric (Fig. 3 and Sup. Fig. 14). The AF predictions were also in very good agreement with the Genesis-trRosetta pipeline with RMSDs of 1.6 to 2.0 Å (Sup. Fig. 14A).

In summary, we showed that our computational pipeline can design various types of folds, including those that have been challenging topologies with beta-rich secondary structures and non-local contacts. Furthermore, the Genesis-trRosetta pipeline was also able to *de novo* design a fold (Top7) that is not found in nature. We utilized a high throughput screening approach to monitor the stability for ∼50 designs for each fold, and through this assay on the order of 10-20 designs out of 50 were resistant to protease digestion suggesting that they are structurally stable or folded. Several designs for each fold were further purified and biochemically characterized, showing that they are folded and stable in solution.

### Exploration of “dark matter” folds

One of the ultimate goals of *de novo* protein design methods is to generate protein folds outside of the natural repertoire potentially amenable to new functionalities. We sought to test whether our framework trained on native protein’s structural data (besides the Top7 fold) is capable of generalizing outside the distribution of natural folds. To this end, we attempted to sample protein folds not included in the training set, nor observed in nature before. Previously, Taylor and colleagues^36^ computationally analyzed unexplored regions of the three-layer mixed alpha/beta fold space through Cα-traces that obey constraints of natural protein structures, such as handedness of SSE and loop connectivity. We further reduced the set by discarding non-designable Cα-traces that had mixed secondary structure types on the same layer, disembodied SSEs, loosely packed or crossing loops. We selected three distinct folds to design with the Genesis-trRosetta method (Fig. 4A).

**Figure 4.**
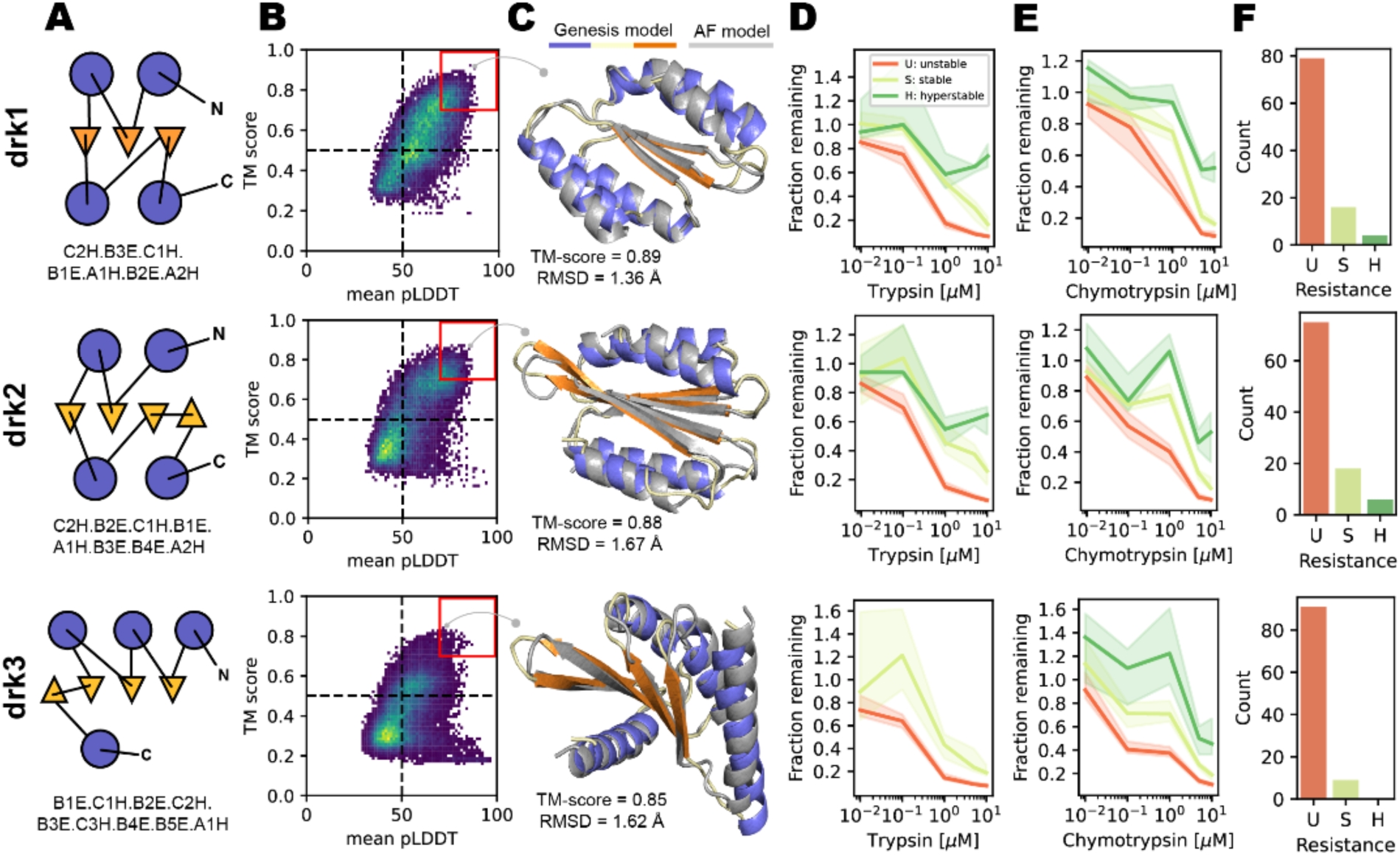
Computational selection and high-throughput experimental stability screening for designed darkfolds. **A:** Forms and 2D representations showing the positioning and connectivity of the SSE elements (circle: helix, triangle: strand) for the three dark-matter folds that guided the fold assembly. **B:** *In silico* assessment of the designs was determined by AF’s mean pLDDT (> 70) and the TM-score between the Genesis and the AF model (> 0.7). **C:** Genesis and AF structural model examples for each fold. **D, E:** Results of an experimental high-throughput enzymatic stability assay on yeast (using Trypsin and Chymotrypsin as enzymes). The libraries were subjected to increasing concentrations of Trypsin and Chymotrypsin proteases, leading to degradation of unstable/unfolded designs. For each design, protease resistance EC_50_s were calculated and the designs categorized as high resistance (H, protease resistance EC_50_ > 5 μM), medium resistant (M, 5 μM > protease resistance EC_50_ > 1 μM), and low resistance (L, protease resistance EC_50_ < 1 μM). The median curve for each stability category is graphed with a confidence interval (CI) of 95 (shaded regions). **F:** Overall summary of the stability measurements of the designs for Trypsin + Chymotrypsin digestions, where designs were required to be either hyperstable or stable with both enzymes.

The first novel fold (drk1) has a central 3-stranded beta-sheet with two helices on both sides. The top helices are connected through the middle and the side strand, and the bottom helices are connected through the other side strand (Fig. 4A top). The second dark fold (drk2) is a three-layer fold with a four stranded beta-sheet in the middle layer and two helices on each side. This fold is similar to drk1, but differs in the connectivity of the top and bottom helices with the central beta sheet (Fig. 4A middle). The third dark fold (drk3) consists of a five-to four-stranded beta-sheet sandwiched with two or three helices on top and a single helix on the bottom. This fold is “rolled” between the four consecutive strands and the three top helices and the last strand connects the lower helix that packs against the full beta-sheet (Fig. 4A bottom).

For the darkfold designs, we followed the same strategy as for the native folds. We used the Genesis-trRosetta framework to sample sequences using different sketches varying in the loop and SSE lengths and collected 2-5 designable candidates (Sup. Fig. 15 and Sup. Fig 16). The selected candidates for each of the three novel folds were then further sampled (Sup. Fig. 16). Structures for all sampled sequences were predicted with AF and the predicted structures compared to the Genesis-trRosetta models. Interestingly, for each of the novel folds a large fraction of the designs exhibited a median TM-score around 0.5 or higher and a median RMSD at 3.7 Å or lower (Fig. 4B). Thus, many of the generated Genesis designs agree with their corresponding AF models (Fig. 4B, C). Based on AF’s pLDDT and the TM-scores, we selected the best 74 designs for further experimental validation. The selected designs for each darkfold were tested in the same manner described above using a yeast display protease-based digestion assay. As for the native folds, a wide range of digestion resistance levels were seen for the designs and each fold had at least one design that present high resistance to protease digestion (Fig. 4D-F). We expressed and purified the top-ranked designs in *E. coli*, and further characterized them biochemically (Fig. 5A).

**Figure 5.**
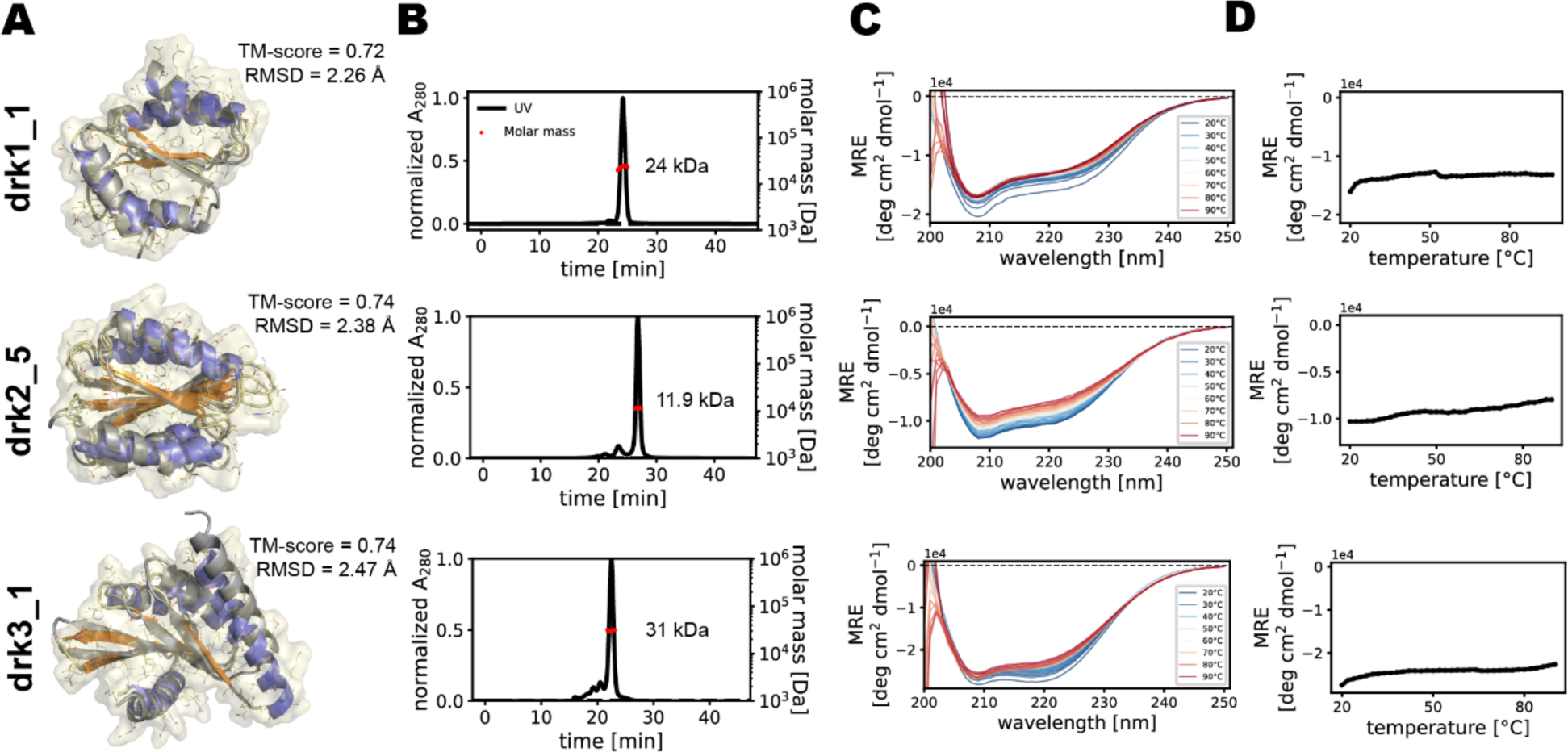
Experimental characterization of the Genesis-trRosetta designs for the dark folds. **A:** Structural model of the top design for each native fold. **B:** SEC-MALS elution profiles showing the oligomeric state of the designs in solution. Red dots show the molecular weight as determined by SEC-MALS. The theoretical molecular weights for a monomer of the designs are: drk1_1: ∼11 kDa, drk2_5: ∼13.7 kDa, drk3_1: ∼13.5 kDa. **C:** Circular dichroism spectroscopy at two different temperatures showing that the designs adopt folded structures in solution. **D:** Thermal denaturation circular dichroism spectroscopy with melting temperatures (T_m_).

For the drk1 fold we attempted 26 designs and obtained 8 that were expressed and purifiable (Sup. Fig. 17). From these, 7 designs presented a monodisperse dimeric species in solution and the expected mixed alpha/beta CD spectra and were thermally very stable, not showing a transition to the unfolded state even at very high temperatures (Fig. 5, Sup. Fig. 17B-D). The RMSDs between the Genesis and AF models varied between 1.9 and 4 Å showing that there was a close agreement between the generated and predicted models (Sup. Fig. 17A). The drk1_1 shown in Fig. 5 is a well-defined dimer in solution, thermally stable and the Genesis model closely resembles the AF model with an RMSD of 2.26 Å.

For the drk2 fold a total of 34 designs were tested, out of which 16 were successfully expressed and purified (Sup. Fig. 18). From the purified designs 4 were present as monomeric in solution and the CD spectra indicated a mixed alpha/beta signature (Sup. Fig. 18B, C). All designs were thermally stable (Sup. Fig. 18D). The AF models agreed with the Genesis models showing RMSDs between 1.9-3 Å (Sup. Fig. 18A). For example, the drk2_5 design that is one of the monomers has a self-consistent RMSD of 2.38 Å (Fig. 5).

Lastly, for drk3 fold we tested 14 designs out of which 3 expressed and were purifiable (Sup. Fig. 19). drk3_3 showed the expected monomeric species in solution and two (drk3_1 and drk3_2) were dimers or small oligomeric species (Fig. 5 and Sup. Fig. 19B). Comparisons between the Genesis-trRosetta models with AF were in very good agreement showing RMSDs between 1.5-2.5 Å (Sup. Fig. 19A).

Taken together, the Genesis-trRosetta pipeline is able to not only generate native folds, but also novel folds not previously sampled by evolution. All the novel darkfolds span across three layers with mixed alpha/beta SSEs. Using a high-throughput enzyme resistance assay on yeast we were able to identify 27 out of 74 designs that were soluble and had the correct secondary structure propensities as measured by CD. This indicates that the Genesis-trRosetta framework can generalize to novel folds not included in the training, suggesting that it has learned structural patterns that are sufficient to guide the backbone generation and sequence search even for folds that have not not been found in nature.

### Mapping successful designs onto the current protein sequence and fold space

The CATH database categorizes protein structure domains into three classes based on their structural features: all-alpha, all-beta, and mixed-alpha/beta. The classes are divided into 40 unique architectures, and architectures are further grouped into 1291 distinct folds. Architectures differ based on the arrangement of SSEs, while folds can have the same architecture but differ in their SSE connectivities.

To evaluate the structural similarity of designed protein sequences to known protein domains, a representative model of each native and darkfold was aligned to all CATH domain representatives using the TM-align^37^ algorithm. The resulting TM-score similarity matrix was mapped onto a 2D plot using classical multidimensional scaling (MDS), revealing that native folds cluster with CATH domains, while darkfolds are separated, suggesting they do not belong to any CATH domain (Fig. 6A, B).

**Figure 6.**
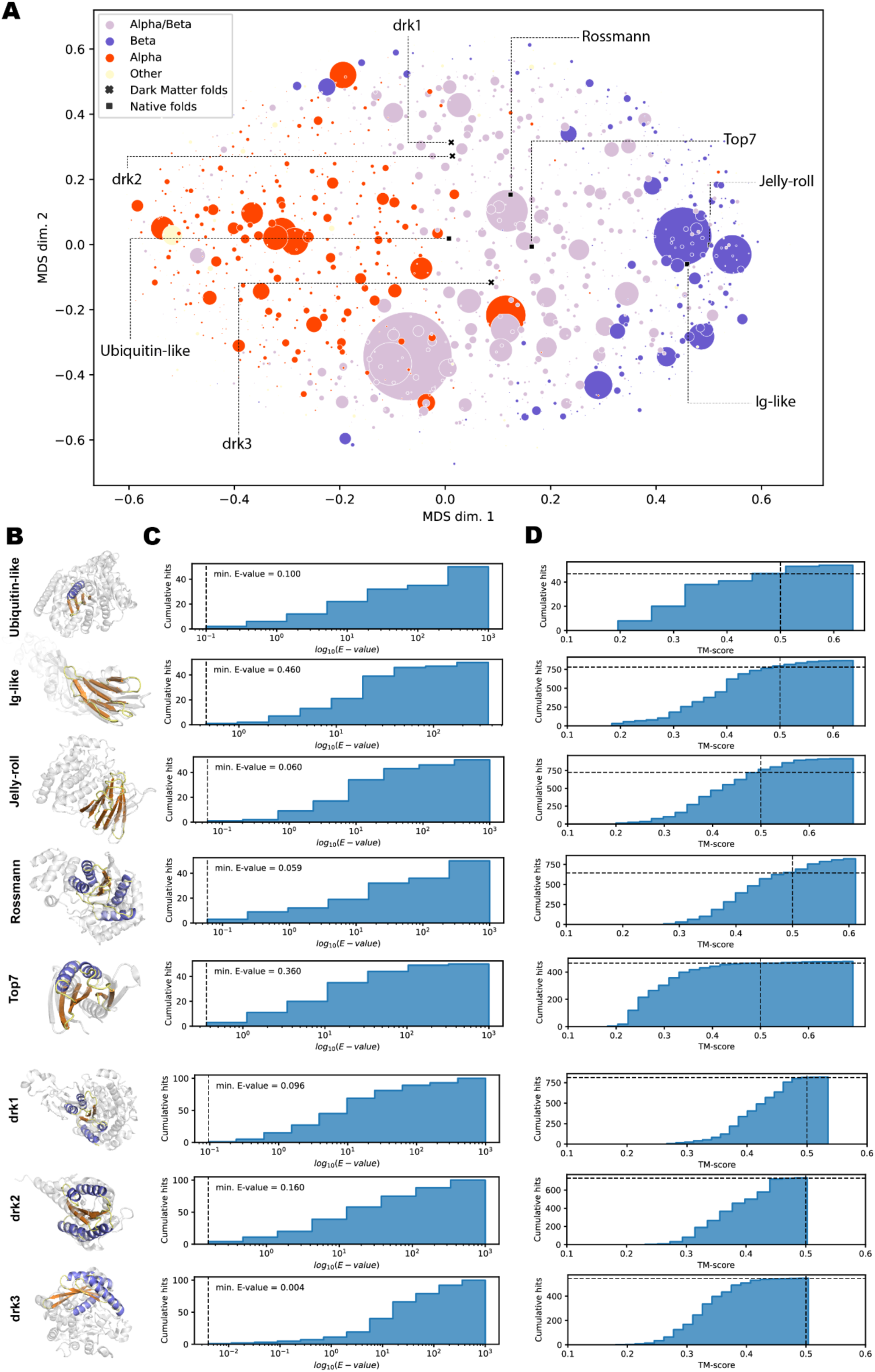
Mapping designs onto the characterized protein space. **A:** Comparison of the protein fold space using TM-scores. The different folds designed are compared using the TM-score to all current folds (by CATH) and mapped via a multidimensional-scaling (MDS). Dark-matter folds are labeled with an **x** and native folds are labeled as ▪. The different sizes of the circles are proportional to the number of members for each of the folds. **B:** Designs mapped to their closest neighbor fold as computed by the MDS. **C:** BLASTp hits against the non-redundant database (NR) of all designed sequences for a particular fold shows that the generated sequences are novel. **D:** TM-alignments for each fold type against the PDB shows that native folds (with exception of Top7) have multiple hits after a TM-score of 0.5, while hits for dark-matter folds are saturated around a TM-score of 0.5 indicating their novelty.

To determine the novelty of the designed sequences, BLASTp^38^ was used to query all designed sequences against the non-redundant database (NR). None of the designed sequences had a hit below the significant E-value threshold of 0.001, indicating that the designed sequences are novel and distinct from any naturally occurring sequence (Fig. 6C).

To further evaluate the closeness of the structural models to native structures, a representative model from each designed fold was used to search the PDB for structural hits using the fast FoldSeek^39^ algorithm (Fig. 6D). Four native folds had multiple matches with a TM-score above 0.5, indicating that these folds effectively align and share the same fold as domains in the PDB. However, the Top7 fold had only a few matches as expected as it is a novel designed fold that did not evolve through time (Fig. 6D). For all three darkfolds, none or very few matches were found with a TM-score close to the 0.5 threshold, indicating that these folds are novel and non-existent in nature (Fig. 6D).

In summary, the designed sequences are novel and distinct from natural sequences. On a structural level, the native designs can be recapitulated within known protein domains, with the exception of Top7, which has a novel fold. The three darkfolds are structurally distinct from naturally occurring domains with TM-scores below or close to 0.5.

## Discussion

We show that a specialized VAE dubbed Genesis is able to encode structural features of low resolution “sketches” of protein folds and decode native-like, designable conformations. By operating on distance-and orientation representations, we are able to (1) alleviate the need of generating designable protein backbones in 3D space with fold-specific restraints and energy functions, and (2) bypass the need for explicit 3D designable backbones in the first place. We coupled Genesis to the trRosetta design engine to generate multiple sequences for the sampled distance-and orientation representations for a set of known and novel folds^20,26^. Our framework is considerably fast, within minutes to generate a sequence and a 3D model for a given target protein shape.

Our results demonstrate that the Genesis-trRosetta framework can design novel proteins that are capable of adopting folds observed in nature and also generalize to novel folds with shapes that are nonexistent in nature. We therefore hypothesize that the Genesis model has learned underlying patterns that govern protein structure. Transforming blurry into sharp patterns, correct protein drafts can drive the trRosetta design method to produce sequences with strong fold signatures. Analysis of the designed protein sequences indicated that they are novel, with most having no close homologs. The use of AF as an orthogonal test demonstrated that a significant proportion of the designed sequences adopt the intended target shape. This indicates that the automated generation of proteins, which were previously only accessible through large-scale simulations, can be enabled by leveraging deep neural networks.

Our experiments suggest that protease-resistant designs were present for both native and darkfold designs, indicating that these proteins are likely to be well-folded. Further comparison of the resistance with that of the native folds from the same family showed that the most stable designs are on par with native proteins. Biochemical analysis of a subset of designs revealed that they adopt well defined species in solution (monomers or low molecular weight oligomers), possess the correct secondary structure composition, and exhibit high thermostability.

Genesis-trRosetta presents a modular approach for designing *de novo* proteins in situations where control over the shape is desired. Our method could be used to create custom protein backbones that conform to non-canonically structured protein interfaces and nanomaterials. Therefore, we anticipate that the versatility and speed of the Genesis-trRosetta method, combined with other possible deep neural network tools for protein design and engineering, can facilitate exploring the vast protein universe and lead to the design of new proteins harboring innovative functions, and overall serve as a valuable resource for future research.

## Material and Methods

### Data set generation

The dataset was built by generating different sets of sketches for large structural tertiary motifs and mapping them to their native counterparts. The sets encompass many 2- and 3-layer fully beta-, fully-alpha-and mixed alpha/beta topological motifs and capture a large scope of possible folds. The sketches used for training have small idealized SSEs (5 AAs residues for strands and 9 AAs residues for helices), no sequence information, and randomly oriented residues along the shortest path between end-and starting points of the secondary structure elements representing placeholders for loops. We created two distinct datasets from the SCOPe (v2.07 stable)^40^ domains of medium sizes (40 AA - 128 AA):

1. **Pretraining dataset with corrupted backbones**: The pretraining dataset was created by corrupting existing protein structures by removing the loops based on the DSSP (hydrogen bond estimation algorithm)^41^ assignments. We remodeled the loops as performed in a sketch, where we added dummy residues (N, C, CA, O backbone atoms with randomized torsion angles) along the shortest path between the two endpoint CA atoms of the consecutive secondary structure elements. We added as many dummy residues as in the native structure, hence the corrupted structure has the same length as its native counterpart. This procedure retains the spatial arrangements of native SSEs. In total, we created a total of 40’726 pairs.
2. **Training dataset with sketches**: We created a set of small sketches obeying simple topological rules such as non-crossing loops and loop distance restraints from the architecture types: EE_EEE, EEE_EEE, H_EEE, H_EEEE, H_EEE_H, HH_EE, HH_EEE, HH_EE_H, HHH, HHH_EE (where “_” represents a layer separation and E: strand and H: helix). We searched SCOPe domains for partial structural matches within 3 Å RMSD using MASTER^42,43^ for each of the generated sketches. These sketches may be either full or partial matches to the native domains. Matching regions of the sketches and native domains are extracted, and SSEs within the extracted structures that do not map to SSEs in the sketch are assigned as loops. oops were modeled this way to avoid biases towards specific conformations. Furthermore, we removed domains larger than 128 AA and identical matches for the mapping to the same sketch. This resulted in a total of 35’435 Sketch - native domain pairs. Despite that sketches can fit onto multiple native counterparts, the majority only maps to 1 or 2 native structures (Sup. Fig. 4).

### Data splits

Within SCOPe^40,44^, protein structures are hierarchically classified into groups where: *Class* groups proteins based on secondary structure content and organization (fully alpha, fully beta, mixed alpha/beta); *Fold* divides structures based on secondary structure element disposition and connectivity; *Superfamily* is based on structural features; *Family* contains the structures with similar sequences. We selected protein *Families* that represent compact structures with small loops for our *Family* test set. The test set includes the SCOPe families b.1.22.1, b.11.1.6, b.69.2.3, b.70.2.1, b.82.1.22, b.114.1.1, a.7.2.0, a.7.2.1, a.7.8.2, a.7.12.1, a.8.11.1, a.24.10.3, a.24.13.1, a.60.9.0, a.160.1.2, c.2.1.7, c.25.1.2, c.118.1.0, c.93.1.0, c.56.5.6, d.110.4.3, e.51.1.1, c.97.1.5, d.17.1.5, d.58.3.2, d.58.10.0, d.58.23.1, d.92.1.13, d.230.1.1, d.240.1.0. Importantly, identical structures and sketches in the training set were removed in order to avoid biases during testing.

### Data encoding

The coordinates of the sketches and their native counterparts were encoded into a total of 4 x 2D distance-and orientation feature maps as done by trRosetta^45^. Briefly, the first feature map represents all-against-all Cbeta distances, the second feature map are dihedral (“omega”) angles that measure the rotation along the virtual axis of two connecting Cbeta residues. The distances and omega angles are symmetric, e.g. measuring from residue 1 to residue 2 will give the same result as measuring from residue 2 to residue 1. The third and fourth features are the “theta” dihedrals and the “phi” angles specifying the direction of Cbeta of residue 2 with respect to residue 1. Both, theta and phi are asymmetric metrics. Together, these 4 feature maps fully define a protein backbone in 3D space.

While we use real valued feature maps as input to Genesis, we bin the true feature maps according to the trRosetta scheme. The distances from 2 Å to 20 Å are binned into 36 equally spaced segments (0.5 Å each) and a 37^th^ bin to indicate that pairs are not in contact. The dihedral (omega, theta) and angular (phi) features are binned into 15° segments yielding 24, 24, and 12 with an additional bin indicating no contact, respectively. Therefore, we have encoded the true feature maps into tensors of shape 128×128×1×37 for the distances, 128×128×1×25 for the dihedrals and 128×128×1×13 for the angles. Thus, at each “pixel” (each residue pair) we have an additional dimension that can be seen as a dirac distribution with a score of 1 for the bin with the distance and 0 everywhere else.

### Genesis architecture

The VAE includes an encoder, a decoder and a loss function. The input Sketch *x* feature maps (real-valued) of shapes 128×128×4 are processed by the encoder, a sequence of 4 convolutional blocks. A single block includes a 2D convolution, an instance norm and an exponential linear unit (ELU) activation followed by a 40% dropout. From the compressed data representation, we use two multilayer perceptrons (MLPs) to predict a normal distribution over the latent space p(**z**|**x**) through predicting a mean *μ* and covariance *σ*^*σ*^ vectors of size 128. Using the reparametrization trick, we sample a latent variable **z** from p(**z**|**x**). The decoder q(**y**|**z**) passes **z** through 3 blocks of 2D deconvolution, instance norm, ELU activation and 40% dropout to create a decompressed representation. The final layer of the decoder branches into 4 different heads. Each head is a convolutional block with a final softmax activation over each pixel yielding distance outputs of shape 128×128×1×37, two dihedral outputs of sizes 128×128×1×25 and an angular output of shape 128×128×1×13.

### Loss function

Our loss function is composed of 5 individual losses (4 reconstruction losses, and a loss on the latent space).

We use the Wasserstein distance^46^ as reconstruction loss. We define **x** ∼ P and **y** ∼ Q and their corresponding densities as p and q, respectively. We assume that (**x,y**)∈ ℝ^*d*^. Additionally, we denote *J*(P,Q) all joint distributions J for (**x,y**) that have marginals P and Q. Then the general Wasserstein distance can be written as:

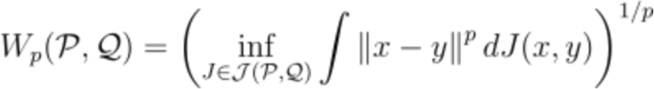

In the discrete case, when P and Q are distributions (x_1, …, x_n) and (y_1, …, y_n) the formulation becomes:

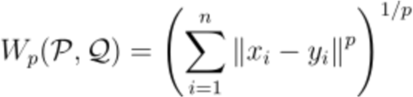

In the case of 1D discrete distributions (p = 1), the 1-Wasserstein (W1) distance is also called Earth mover’s distance (EMD) and is efficiently computed. The main advantage of the 1-Wasserstein distance compared to other measures, such as the binary cross-entropy and the Kullback-Leibler (KL) divergence, is that it takes into account the metric space. This means that larger deviations from the predicted to the true distributions are more penalized while small errors are less penalized.

We define the reconstruction loss (*L_rec_*) as the sum over the 1-Wasserstein distances between the predicted distributions (*Dhat_p_*) and the true distributions (*D_p_*) of each pixel normalized by the length of the protein (*N_AA_*). Each pixel is defined as (i,j) where i = 1,…,n_w and j=1,…,n_h with n_w being the width and n_h the height.

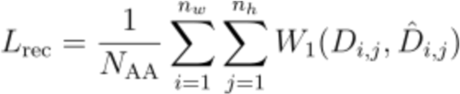

Note that the true distribution is modeled as a Dirac distribution supported by the true values, whereas the predicted distribution (*Dhat_p_*) is parametrized by the VAE decoder.

We additionally use the Kullback-Leibler (KL) divergence on the latent space normalized by the length of the protein to penalize latent vectors not following a Normal distribution KLD = 1/N_AA_ * KL(p(z|x) || p(z)), with p(z) ∼ Normal(0,1). Thus the final loss is defined as:

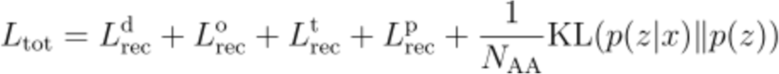

where d, o, t and p are the distances, omegas, thetas and phis, respectively.

### Training regimen

We followed a standard pretrain - fine-tune regimen. We pretrained the VAE on the corrupted structures with a learning rate set to 1e-3 over 300 epochs and subsequently fine-tuned for 500 epochs on the sketches. We reduce the learning rate with a step of 0.97 at each epoch during pre-training and every second epoch during fine-tuning. A batch size of 64 sketches was used with the Adam optimizer^47^. The pretraining slightly improved the performance on the test set when compared to directly training the VAE on the sketches.

We trained the VAE in a supervised manner by minimizing the 1st Wasserstein distance between the true feature maps and the distribution predicted by the VAE. In contrast to the previously utilized cross-entropy loss, the Wasserstein distance enables weighting individual errors between the distributions, i.e., penalizing large differences between the true and predicted distributions more than small differences.

### trRosetta design engine and modeling

The trRosetta design framework was utilized to design a set of 1K sequences matching Genesis refined inter-residue distance and orientation maps. A position-specific scoring matrix (PSSM) was generated from the library of sequences and used within the PyRosetta^48^ protocol. In a first stage, the PyRosetta protocol generates a coarse-grained model using gradient descent with the optimized restraints and a single sequence from trRosetta design. In a second stage we removed the restraints assuming that the generated coarse grain model has adopted the target shape. We further optimize the coarse grain structure with a full-atom protocol. We use the Rosetta FastDesignMover with layer and PSSM sequence constraints during the design task and topological secondary structure energy bonuses during the relaxation task. In this way, the full-atom protocol improves the quality of the final sequence and structure model.

### Candidate stage generation

The candidate search stage samples different Forms in order to identify designable SSE-and loop length combinations (SSE and loop lengths that fit together). To evaluate the feasibility of a Form in 3D, the Genesis-trRosetta framework was used to generate protein models that align with a TM-score of 0.5 or higher onto the input sketch. Here, we gave the Genesis-trRosetta framework two attempts to fulfill the TM-score criteria. Note that the number of attempts can be increased for a more fine-grained search. Generally, it was assumed that a 3D model with the correct connectivity would have a TM-score above 0.5 to the sketch, indicating that it has the same fold. For the Rossmann-, Jellyroll-and Top7 folds, more than 50% of the candidates passed the 0.5 TM-score threshold and were collected, while for the Ig-like and Ubiquitin-like folds, approximately 25% passed the TM-score threshold.

### Production stage generation

Each successful candidate from the search stage was subjected to a large production design generation. Specifically, the Genesis-trRosetta framework was used to design up to 20’000 sequences for a specific designable sketch. For all designs, we used AF to predict the structural models of the designed sequences. A substantial number of designed sequences for all folds are accurately predicted by AF as assessed by TM and confidence (pLDDT) scores. We selected the top 250 for native folds and top 250 sequences for the darkfolds by TM-scores and pLDDT for further experimental analysis.

### Protein expression and purification

DNA sequences of the designs were purchased from Twist Bioscience as fragments or from Genscript as cloned plasmids. The DNA fragments were cloned via Gibson cloning into a pET21b followed by a terminal His-tag. All plasmids were sequenced and transformed into Escherichia coli BL21(DE3). Expression was conducted in Terrific Broth supplemented with ampicillin (100 μg/mL). Cultures were inoculated at an OD600 of 0.1 from an overnight culture and incubated in a shaker at 37 ℃ and 220 rpm. After reaching an OD600 of 0.6-0.8, expression was induced by the addition of 1 mM IPTG and cells were further incubated overnight at 18 ℃. Cells were harvested by centrifugation and pellets were resuspended in lysis buffer (50 mM TRIS, pH 7.5, 500 mM NaCl, 5% glycerol, 1 mg/ mL lysozyme, 1 mM PMSF, 4 μg/ml DNase, 0.5X Cell Lytic lysis reagent). Resuspended cells were incubated with shaking for one to two hours and clarified by centrifugation. Ni-NTA purification of sterile-filtered (0.22 μm) supernatant was performed using a 5 ml His-Trap FF column on an ÄKTA pure system (GE Healthcare). Bound proteins were eluted using an imidazole concentration of 300 mM. Concentrated proteins were further purified by size exclusion chromatography on a Hiload 16/600 Superdex 75 pg column (GE Healthcare) using PBS buffer (pH 7.4) as mobile phase. The peak corresponding to the expected molecular weight was collected and concentrated for analysis.

### Circular dichroism spectroscopy

Far-UV circular dichroism spectra were collected between wavelengths of 200 and 250 nm on a Chirascan™ spectrometer (AppliedPhotophysics) spectrometer in a 1 mm path-length quartz cuvette. Proteins were prepared in PBS at a concentration of 0.3 mg/mL (or less if precipitating at 0.3 mg/mL). Wavelength spectra were averaged from two scans with a scanning speed of 20 nm/min and a response time of 0.125 s and each measurement was reference subtracted using a cuvette with PBS. The thermal denaturation curves were collected by measuring the change in ellipticity at 220 nm from 20 to 90 ℃ with 2 ℃ increments.

### Size-exclusion chromatography combined with multi-angle light scattering

Multi-angle light scattering was used to assess the monodispersity and molecular weight of the designed proteins. Samples containing 50–100 μg of protein in 100 uL PBS buffer (pH 7.4) were injected into a Superdex 75 10/300 GL column (GE Healthcare) at a flow rate of 0.5 ml per minute coupled in-line to a multi-angle light-scattering device (DAWN TREOS, Wyatt). Static light-scattering signals were recorded and the scatter data were analyzed by ASTRA software (version 8.0.2.5 64-bit, Wyatt).

### Yeast library preparation

Designs were ordered as gene fragments from Twist Bioscience and amplified individually (to avoid PCR bias for certain designs) using pNT_homo_fwd and pNT_homo_rev primers (see Table S4.1). After amplification, the reaction mixtures were pooled and purified. The pNT-V5 display vector was digested with NheI, BamHI, and SalI and purified by column for electroporation. Oligo pool libraries were ordered from Twist Bioscience. The pools were amplified using minimal cycles using pNT_min_fwd and pNT_min_rev primers for the first round, and pNT_min_fwd_2 and pNT_min_rev_2 for the second round. For both pools and individual designs, the insert and backbone were combined in a mass ratio of 5:1 and the DNA dehydrated for transformation.

For the yeast library transformation, a protocol published in Chao et al^49^ was followed. Briefly, an overnight culture of EBY100 in YPD containing penicillin streptomycin (pen/strep) was used to inoculate a 100 mL culture at an OD600 of 0.1. The culture was grown at 30℃ with 200 rpm shaking, until the OD600 reached 1.3 (about 6 hours). 1 mL of 2.5 M DTT (freshly prepared in 1 M Tris, pH 7.5) was added for a final concentration of 25 mM and the flask was returned to the shaker for 10-15 more minutes. The culture was pelleted in two pre-chilled tubes by centrifugation at 4000×g for 5 minutes at 4℃. The pellets were rinsed in 25 mL E buffer (10 mM Tris, pH 7.5, 270 mM sucrose, 2 mM MgCl_2_), repelleted, and rinsed in 1 mL of E buffer. The suspension was transferred to a 1.5 mL Eppendorf tube, pelleted, and both pellets resuspended in 300 uL E buffer total. The yeast slurry was added to the dried DNA tubes (40 μL of slurry per cuvette), incubated on ice for 10 minutes, and electroporated (with settings of 0.54 kV and 25 μF and no pulse controller). After electroporation, the yeast were recovered in YPD media for one hour before pelleting and performing serial dilutions to check transformation efficiency. The rest of the culture was added to 100 mL SDCAA media with pen/strep and grown overnight.

### Digestion and sorting of yeast libraries

Yeast libraries were passaged at least once before inducing for 18 hours at 30℃ in SGCAA media with pen/strep. Induced yeast were pelleted by centrifugation at 3000×g for 3 minutes at 4℃. The amount of yeast pelleted corresponds to 2 mL at an OD600 of 1 for each digestion sample. The pellets were resuspended in either PBS (pH 7.4, for trypsin digestion) or TBS (20 mM Tris pH 8.0, 150 mM NaCal, for chymotrypsin digestion). After rinsing once, the pellets were resuspended in a volume of 250 μL per sample and aliquoted into tubes for digestion.

The trypsin and chymotrypsin were freshly resuspended from powdered stocks of enzyme and used within one hour of resuspension. Both were prepared as 2x concentrations and 250 uL added to each tube. The final concentrations used were 0.01 μM, 0.1 μM, 1 μM, 5 μM, and 10 μM. After mixing with the yeast, the digestion was allowed to occur for 5 minutes before quenching with ice cold 2% BSA in PBS. The yeast was immediately pelleted and rinsed three times with 2% BSA in PBS. After the final rinse, the supernatant was carefully removed and 50 μL of 0.1% BSA in PBS with 1:100 anti-HA-FITC (clone GG8-1F3.3.1). The tubes were rotated end-over-end for 15- 30 minutes in a foil wrapped tube. The yeast was pelleted and the sample resuspended in 0.5 mL of 0.1% BSA in PBS for sorting.

The genetic construct used for display is a modified version of pNT vector that encodes an HA epitope tag, the protein of interest, a V5 epitope tag, and then the Aga2 protein (Sup. Fig. 9). To test stability, the loss of the HA epitope signal is used as a proxy for protein degradation. The construct displaying a stable protein is able to withstand up to 10 μM trypsin and chymotrypsin without a significant drop of signal (Sup. Fig. 9), thus any drop in signal in our assay can likely be attributed to degradation of the designs.

Sorting was performed using a SONY SH800 instrument. To set the gates, an unlabeled sample and a labeled sample with no digestion were run and a histogram of number of events versus FITC-fluorescence generated. The positive gate was drawn such that no more than 1% of the unlabeled sample events were occurring in this gate. After setting the gate, it was not altered for any samples in the sorting set. For each sample, the sorter was set to Ultra-purity mode and 5 million events were allowed to occur while collecting cells from the positive gate into SDCAA. After the sort, SDCAA with pen/strep was added to each sample and all were shaken overnight (or up to three days) until a substantial density was reached.

### MiSeq sample preparation and sequencing

Sorted yeast were passaged once into 2 mL of SDCAA media with pen/strep and grown to an OD of about 1. The yeast were pelleted and the DNA purified using Zymoprep Yeast Plasmid Miniprep II kit. The instructions were followed except for additional steps of flash freezing and thawing before addition of the Zymolase and after the incubation at 37℃ degrees, to increase the yield. To prepare the DNA for MiSeq sequencing, primers listed in table S4.1 were used for PCR I (details is Table S4.2) to copy out the sequences of interest and add the adapters for the Illumina sequencing barcodes. The products were purified with Qiagen quick clean up kit. Standard Illumina sequencing primers were used for PCR II (details in Table S4.2) to add barcodes, with a unique set for each of the samples. PCR II products were also purified, with 2 extra washes of the column to ensure the final sample had minimal salt contamination. The DNA was eluted with water and the concentration quantified with a Qubit dsDNA high sensitivity kit. The PCR II products were run on a Fragment Analyzer to determine if the majority of the DNA present was the expected size and ensure the absence of primer-dimers.

Finally, samples were analyzed with a MiSeq benchtop sequencer by the Gene Expression Core Facility at EPFL, using standard protocols. 500 paired end cycles were run, with approximately 1 million reads per sample.

### Protease resistance determination for libraries

The FASTQ files from the MiSeq experiment were processed using python. First, files were trimmed and filtered to contain lists of only sequences that start and end with the desired sequences and would be in frame. Then, the sequences were translated into amino acid sequences. The sequences of the designs included in the library were tabulated to count the number of instances of each sequence in each sample. This table was used to normalize the amount of yeast displaying each design in each sample.

The relative amount of each design was normalized to 1 and generally decreased, as expected. Each set of points was fit to an IC_50_ curve with fixed top and bottom (1 and 0, respectively). The designs were then given an IC_50_, referred to in the text as a folding score. For very resistant designs, the relative amount of yeast displaying that design does not decrease and the fitting gives a large number, which is listed as >10 μM, as this was the highest concentration of enzyme in our assay.

### Protease resistance analysis for single designs

Genes encoding native sequences for each of the 5 known folds were ordered from Twist Biosciences with flanking regions for pNT yeast display vector. PCR was performed with the designs as in Supplementary Table 2. The inserts were transformed into yeast using Zymoprep yeast transformation kit II with cut pNT V5 vector. Individual designs were grown and induced in the same manner as the libraries. Additionally, 2 or 3 Genesis-generated designs from each fold were included to benchmark the native sequences with the designs and compare to the protease resistance for libraries.

To test the protease resistance for the individual proteins, induced yeast were digested with various concentrations of trypsin and chymotrypsin (from 0.01 μM to 10 μM in ten-fold steps) for five minutes at room temperature. After quenching and washing three times with ice cold PBS supplemented with 1% BSA, the samples were incubated with anti-HA FITC antibody. An Attune NxT was used to read 10,000 cells from each sample.

The percentage of FITC positive cells in the labeled, undigested sample was used to normalize the FITC-positive population for each of the digested samples. As with the library samples, Prism was used to fit a curve and calculate a resistance score for each protein. Supplementary table 2 lists the calculated resistance score for each protein testing. The Genesis-designed proteins that were included as benchmarks indicate the resistance score from both the individual test and the library test.

### Protein BLAST analysis

The design sequences were run through BLASTp^38^ to compare against the non-redundant protein database (all non-redundant GenBank CDS translations+PDB+SwissProt+PIR+PRF excluding environmental samples from WGS projects) with 512,512,737 sequences. The expected values (E-values) returned by BLASTp indicate the number of times a match of the given quality might be “expected” to be found by random chance within the database searched. For the search, the maximum E-value was increased to 1000. The E-value of the closest match was tabulated for each sequence. Any sequences that returned no matches were listed as having an E-value of 1000.

## Acknowledgments

We thank the members of the Protein Design and Immunoengineering group (LPDI, EPFL, Lausanne) for helpful discussions. We thank Martin Pacesa for critical reading of the manuscript. We would like to thank EPFL’s Scientific IT and Application Support Center for their support on the computational infrastructure. We would like to thank the Protein Production and Structure Core facility at EPFL for their support on the protein biophysical characterization experiments. B.E.C. is a grantee from the European Research Council (Starting grant - 716058), the Swiss National Science Foundation, and the Biltema Foundation. Parts of the computational simulations were performed at the CSCS - Swiss National Supercomputing Centre through a grant obtained by B.E.C.. Z.H., and S.R. are supported by a grant from the National Center of Competence in Research in Chemical Biology. JS is supported by the UKRI CDT in AI for Healthcare http://ai4health.io (Grant No. P/S023283/1). M.M.B acknowledges the European Research Council Consolidator grant 724228 (LEMAN). We thank Willie Taylor for providing the initial models for the dark folds.

## Data availability

The Genesis code and trained weights can be downloaded at https://github.com/zanderharteveld/genesis

## Additional information

### Author information

These authors contributed equally: Zander Harteveld, Alexandra Van Hall-Beauvais.

### Contributions

B.E.C. and Z.H. conceived the initial idea and refined it together with Z.H., A.L, M.D., J.S., P.V., M.M.B. and B.E.C.. Z.H. designed and performed experiments. Z.H. wrote the software with support from J.S. and A.L.. A.K.V., S.G., I.M. and S.R. purified the designs and performed protein-biochemical characterization.. A.K.V. performed the yeast library preparation, sorting, and DNA preparation for MiSeq. A.K.V. and Z.H. analyzed the MiSeq data with assistance from A.Z.. Z.H. performed *in silico* structural analysis and modeling. C.G. performed sequence analysis using pBLAST. J.S. performed structural comparison of designs against the CATH database. B.E.C. directed the work. Z.H., A.K.V. and B.E.C. wrote the manuscript with support from all authors.

