## Supplementary material for "Exploring “dark matter” protein folds using deep learning": Supp material

### Supplementary Figures

**Figures**:

- Overview of VAE
- Sketch - native comparison
- General sampling problem
- Genesis exploration - exploitation strategy
- Native fold length and candidate metrics
- Darkfold length and candidate metrics
- Configurations (Forms) sampled
- Genesis denoised feature maps for native folds
- Genesis denoised feature maps for darkfolds
- Display construct and gating
- Biochemical data compared to native folds
- Biochemical data for native folds
- Biochemical data for darkfolds


**Tables**

- Sequences for tested designs

**Sup. Figure 1.**

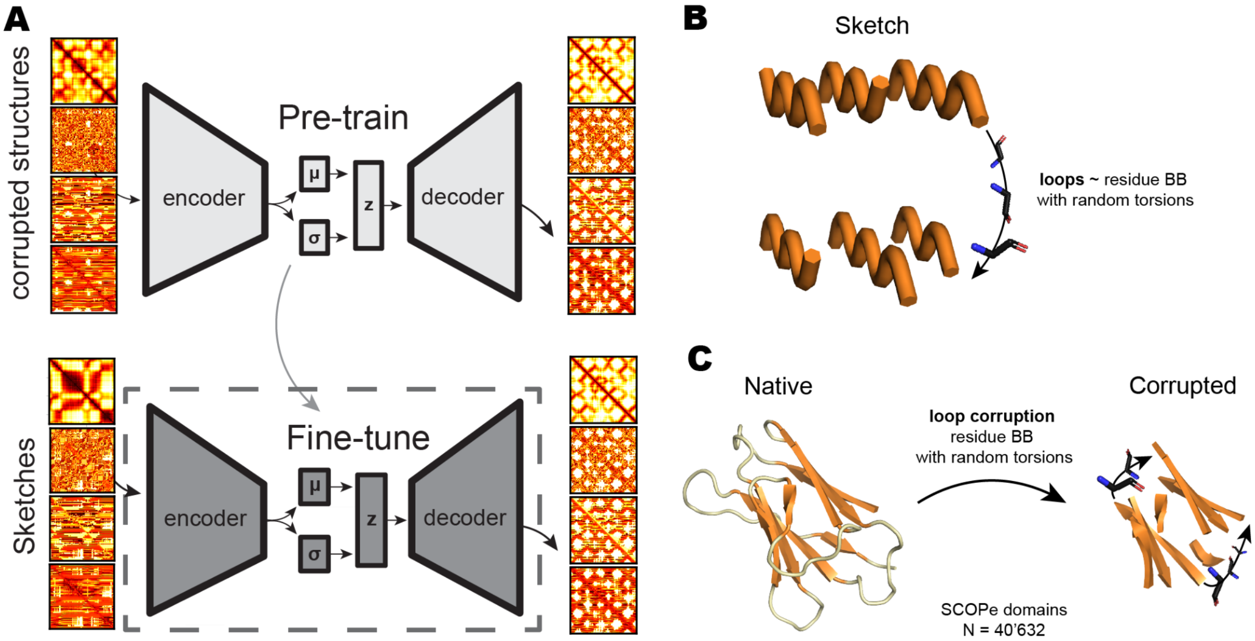


**Sup. Fig. 1 | Overview of VAE architecture, training regimen and loop representation.**

**A:** Overview of the training process for Genesis. The VAE was first trained on a large dataset with loop-corrupted structures and then fine-tuned on a smaller dataset with sketches. **B:** Loops in the sketches are represented as randomly rotated disconnected backbone residues along the shortest path connecting idealized SSEs start and endpoints. **C:** Similarly, loops in the corrupted structures are represented as randomly rotated disconnected backbone residues along the shortest path connecting SSEs start and endpoints.

#### Sup. Figure 2.

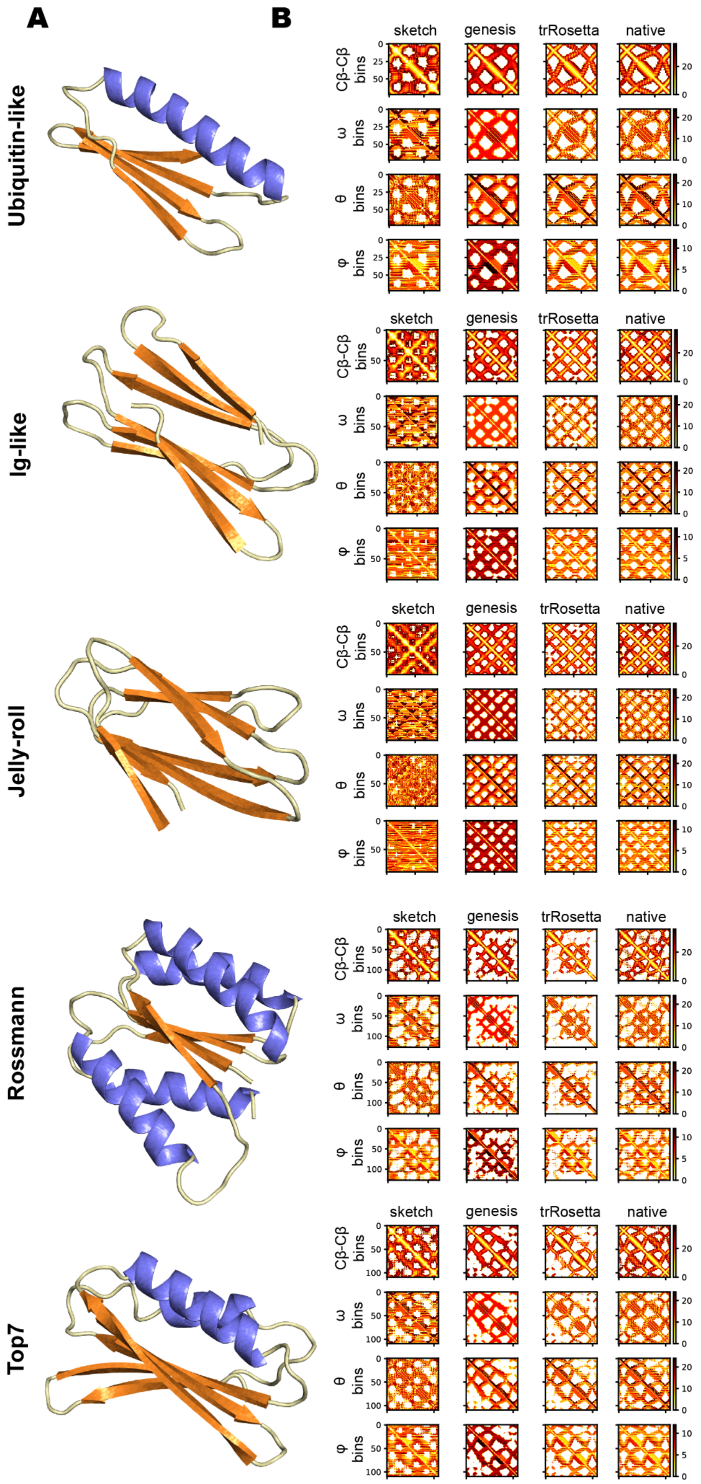


**Sup. Fig. 2 | Examples of Genesis-optimized distances for native folds.**

**A:** Examples of structural models from Genesis for each fold. **B:** Features distance- and orientograms of various stages during the Genesis-trRosetta protocol showing gradual improvements and denoised representations indicating improved designability.

**Sup. Figure 3.**

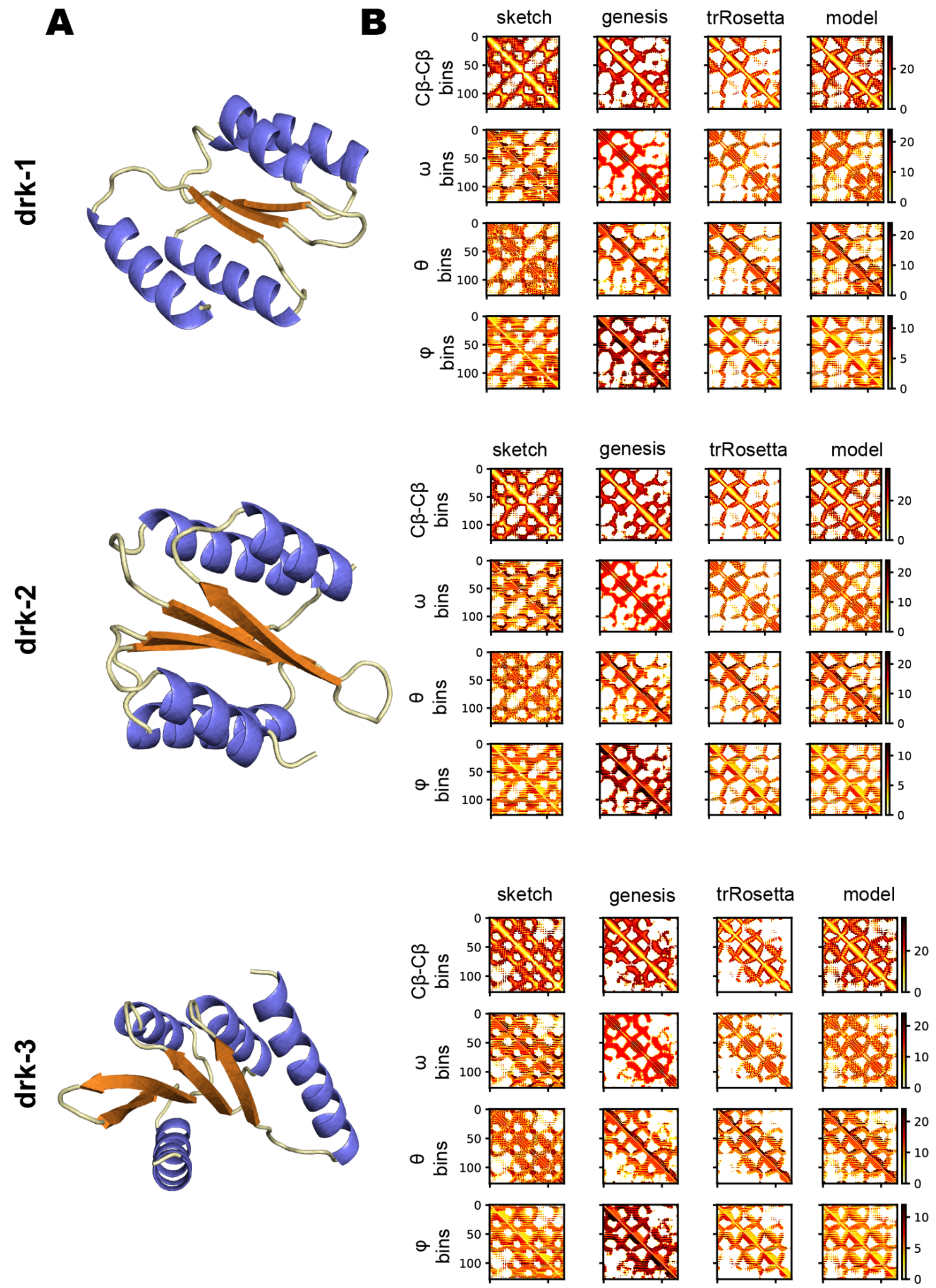


**Sup. Fig. 3 | Examples of Genesis-optimized distances for darkfolds.**

**A:** Examples of structural models from Genesis for each darkfold. **B:** Features distance- and orientograms of various stages during the Genesis-trRosetta protocol showing gradual improvements and denoised representations indicating improved designability.

**Sup. Figure 4.**

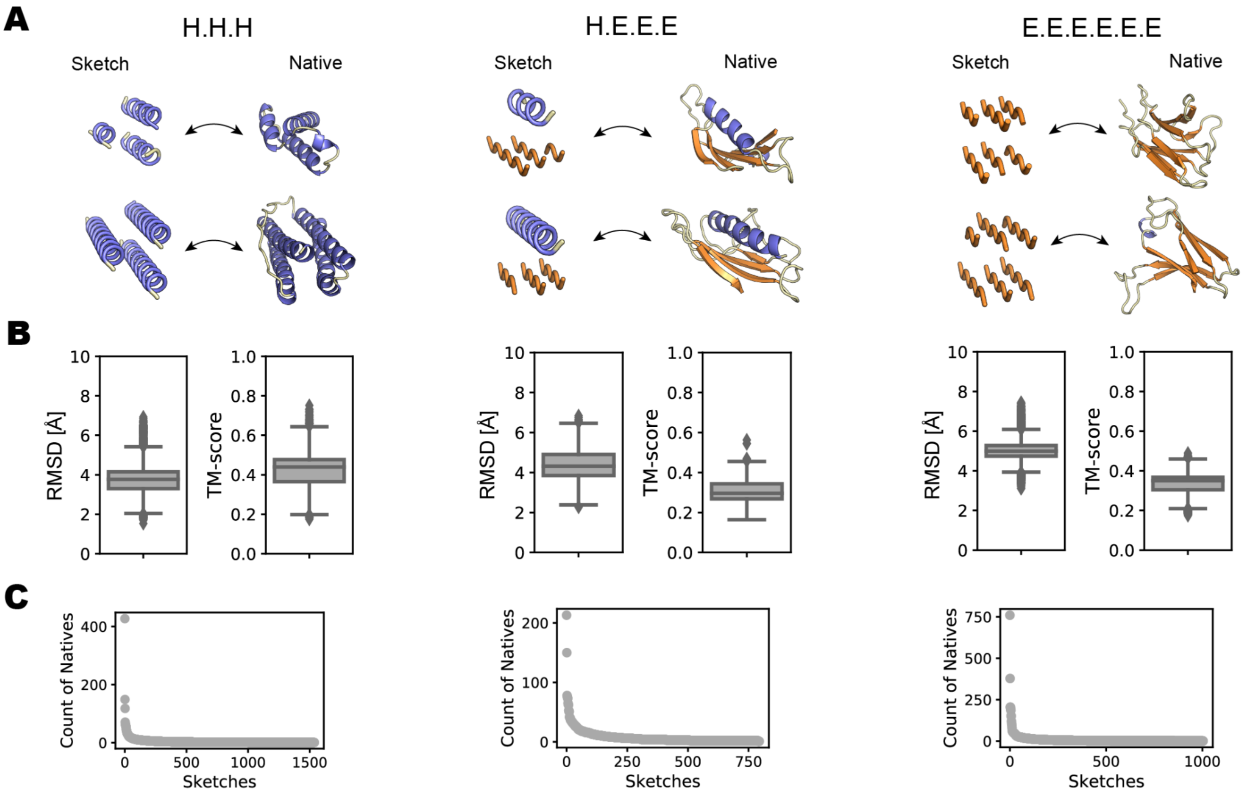


**Sup. Fig. 4 | Comparison of protein fold sketches and their corresponding native structures.**

**A:** Examples of sketches and their corresponding native structure counterparts. **B:** Alignments of the sketches onto their native matches shows that the sketches are oftentimes very different and lack structural similarity. **C:** Number of native backbones that fit onto a given sketch (RMSD <2 Å). Multiple native counterparts can fit onto one sketch, but most sketches have only a few native conformations.

#### Sup. Figure 5.

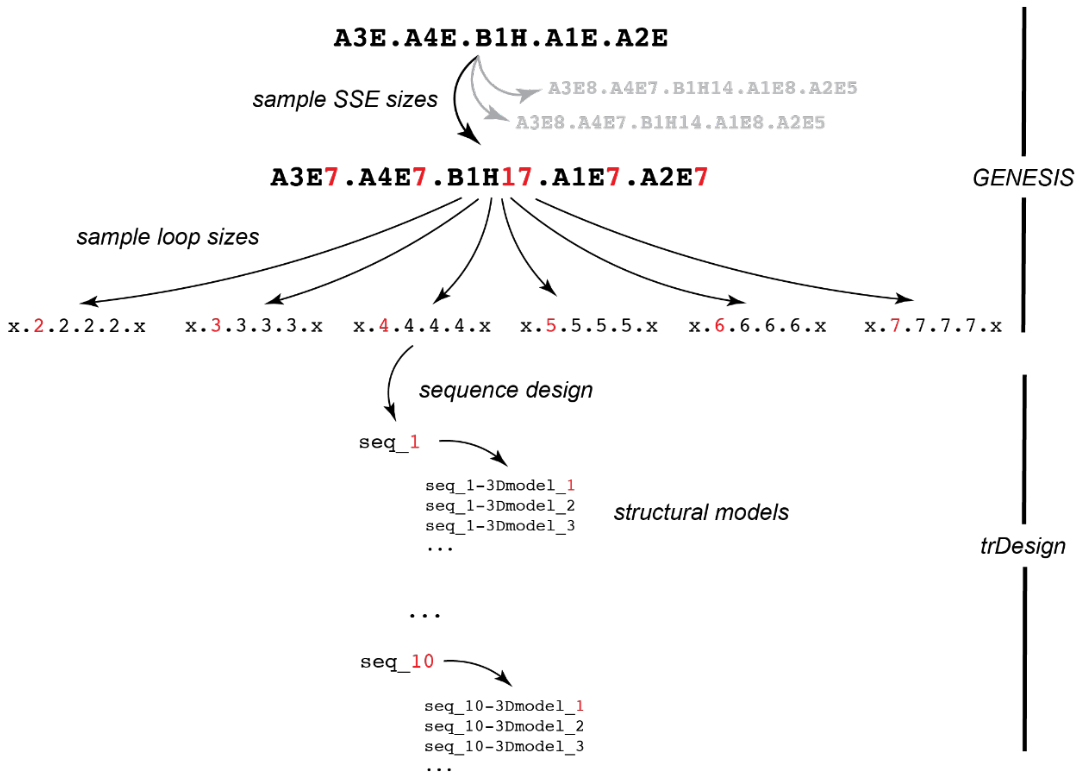


**Sup. Fig. 5 | General overview of the sampling strategy for the Genesis-trRosetta pipeline.**
Overview of the sampling strategy employed for the Genesis-trRosetta pipeline indicating the choices and levels at which the modules were used. A sketch describing a layered fold is provided as input, followed by (1) sampling over several SSE length configurations and (2) loop length combinations. For each of the sketches with different SSE-loop lengths combinations, (3) several sequences were then designed using trRosetta in independent trajectories and (4) several 3D models were assembled via PyRosetta for each designed sequence.

#### Sup. Figure 6.

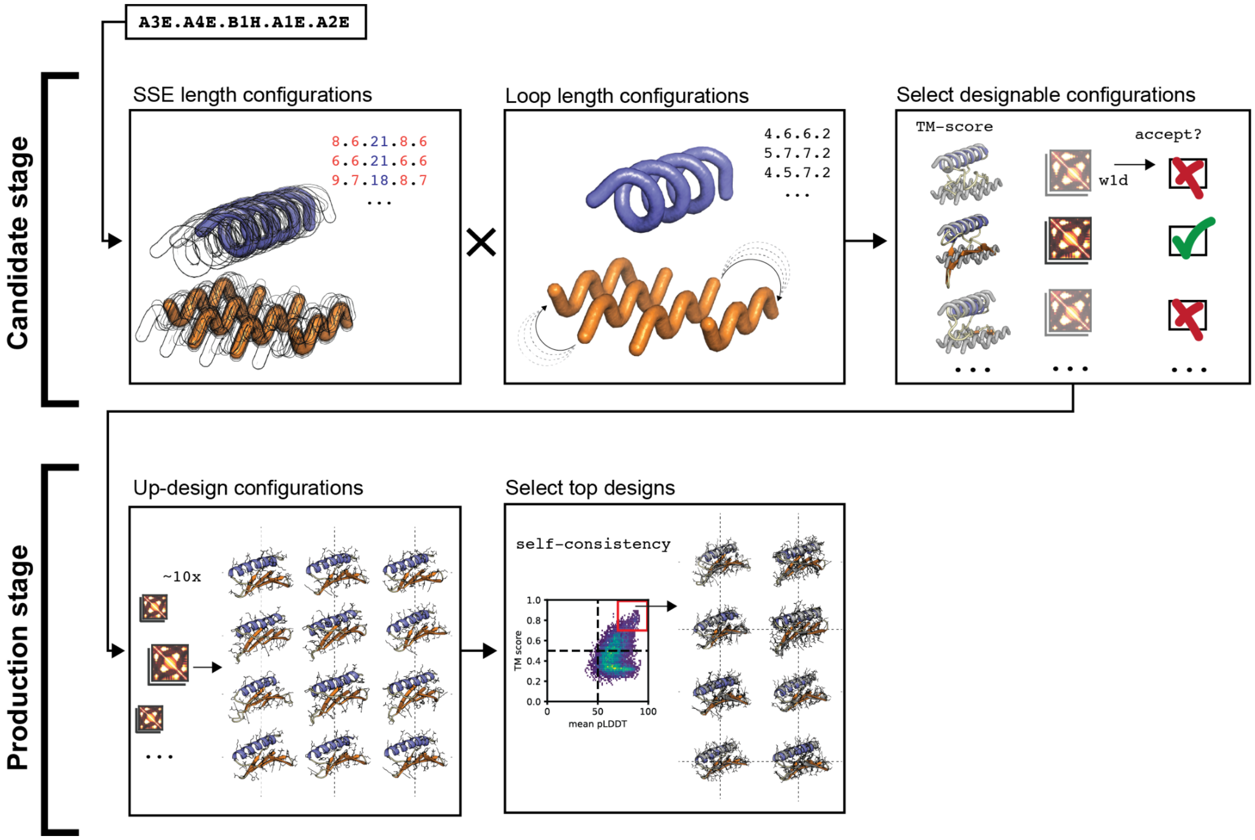


**Sup. Fig. 6 | Exploration and exploitation sampling strategy using Genesis**

The exploration sampling involves searching for designable candidates: (1) the fold is defined using a form string description specifying the location of the SSE in 3D space; (2) a sketch is created from the form, SSE lengths are sampled; (3) for each SSE-length combination different loop sizes are approximated. This is achieved using an algorithm that first computes the minimum number of residues needed, and then randomly returns loop size combinations within the minimum length and up to + 3; (4) the SSE-loop-size combinations are generated and designable candidates selected based on the TM-score between the Genesis output and the Sketch (> 0.5 indicating the correct fold) and the average 1st Wassterstein distance between the Genesis-trRosetta and Genesis-model maps (indicating self-consistency between structural backbone features). The production stage involves exploiting the designable candidates which includes: (1) designing a large pool of sequences from selected Genesis features maps, and (2) selecting the top sequences based on TM-score between the Genesis and AF models, and the AF pLDDT score (> 70).

**Sup. Figure 7.**

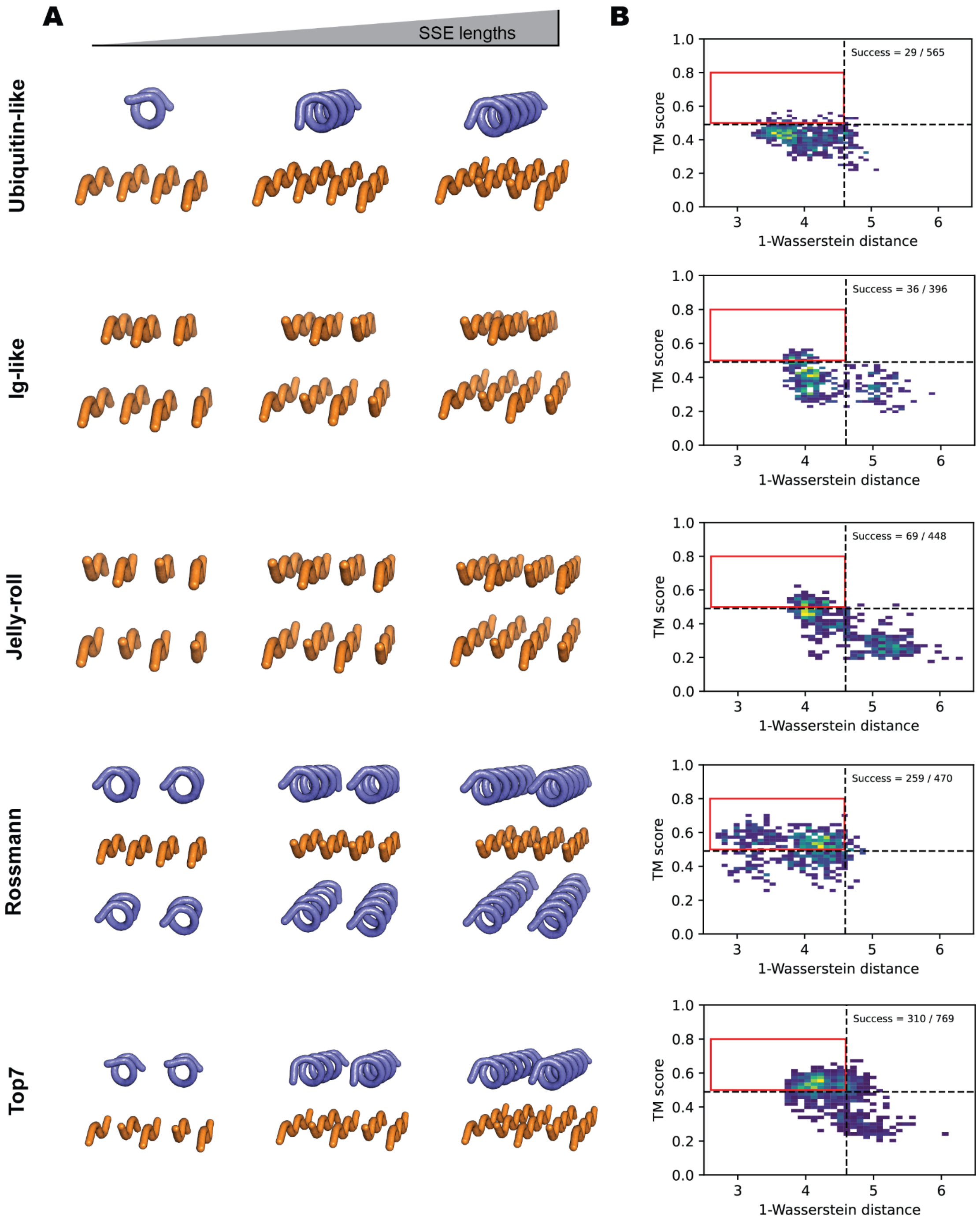


**Sup. Fig. 7 | SSE size configurations for native folds span a wide range of lengths.**

**A:** Examples of different SSE length combinations sampled during the exploration stage to find designable candidates. As shown, a wide range from small to large SSEs were searched for potential designability. **B:** Accessing backbone designability of candidates. First, the TM-score between the Genesis model and the input Sketch (> 0.5) indicates if the generated model retains the correct fold and the averaged 1st Wasserstein distance between the Genesis and trRosetta feature maps, and Genesis model feature maps indicating consistency between denoised features through the design stages.

**Sup. Figure 8.**

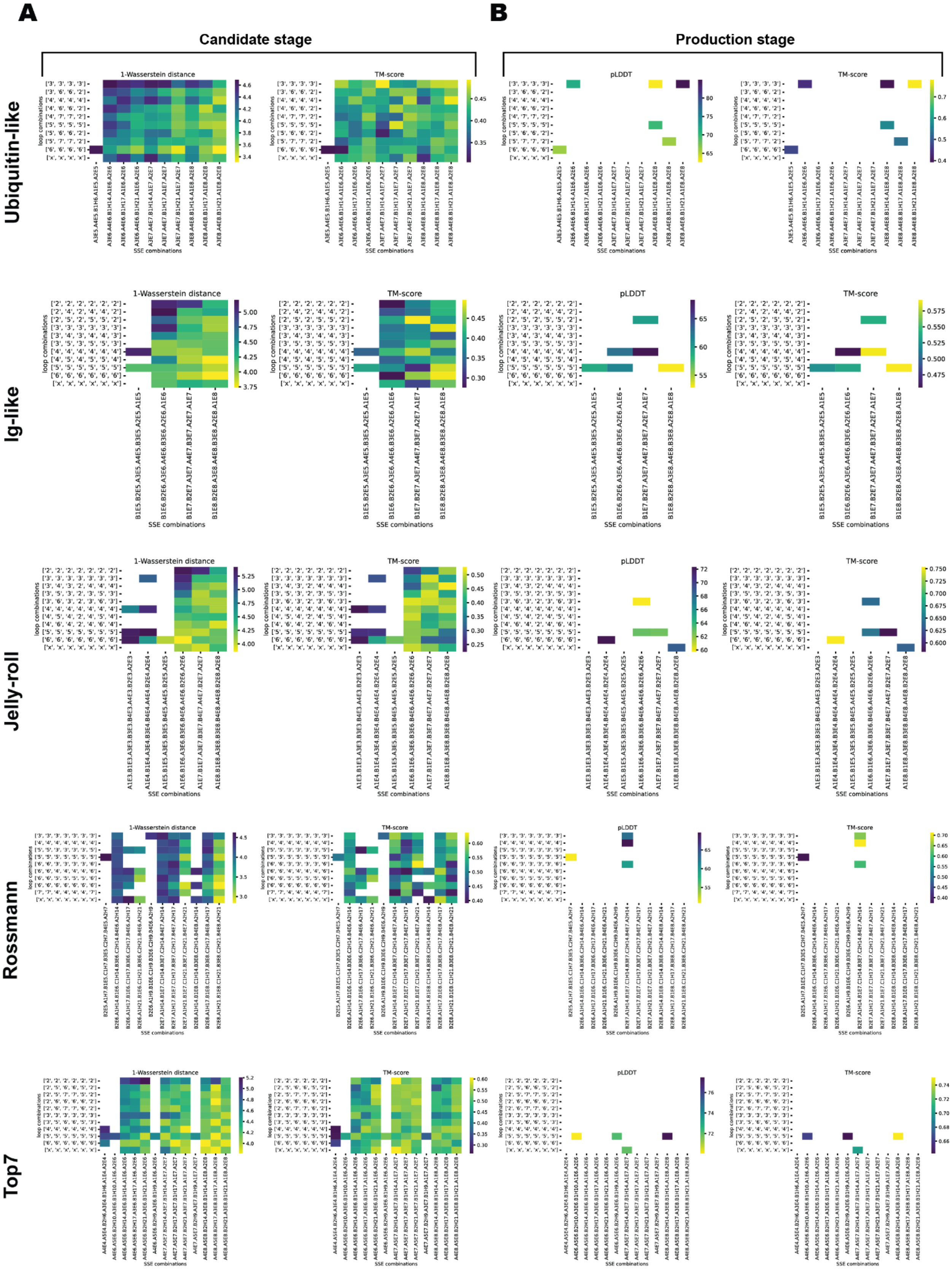


**Sup. Fig. 8 | Score-based selection of combinations for candidate and production stages for native folds.**

**A:** Computational scores (1-Wasserstein distance and TM-score metric) of designs for different loop and SSE combinations sampled at the candidate stage. Candidates with a 1-Wasserstein distance below 4.6 and a TM-score above 0.5 were considered for the production stage. **B:** Computational scores (pLDDT and TM-score) of upsampled designs from the candidate stage used as selection criteria.

**Sup. Figure 9.**

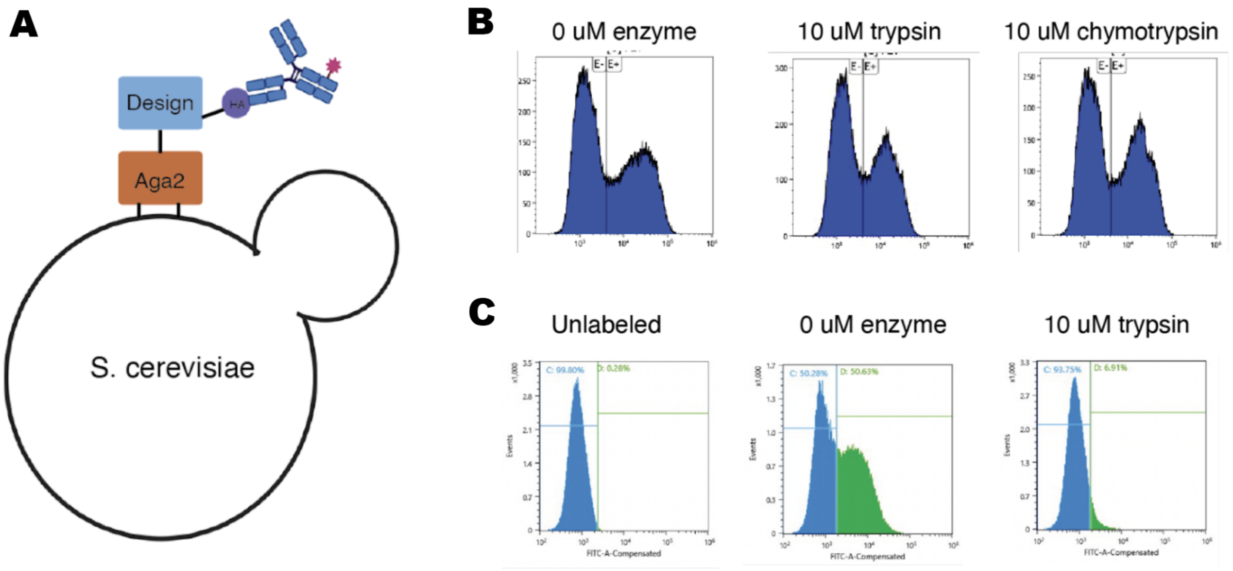


**Sup. Fig. 9 | Display constructs and gating.**

**A:** The yeast display construct displays, from N-terminus to C-terminus, an HA-epitope tag, the designs, and the Aga2 protein. An anti-HA-FITC antibody is used for detection of full length constructs. **B:** Testing the stability of the construct to trypsin and chymotrypsin. A stable, folded design is displayed on yeast and subjected to 10 μM of protease. If the construct retains the display tag this indicates that the display construct itself is stable. **C:** The gates for sorting were set using unlabeled yeast (left) and undigested yeast (center). As the enzyme concentration increased, the amount of FITC-positive cells decreased, but the gate remained constant for all samples.

#### Sup. Figure 10.

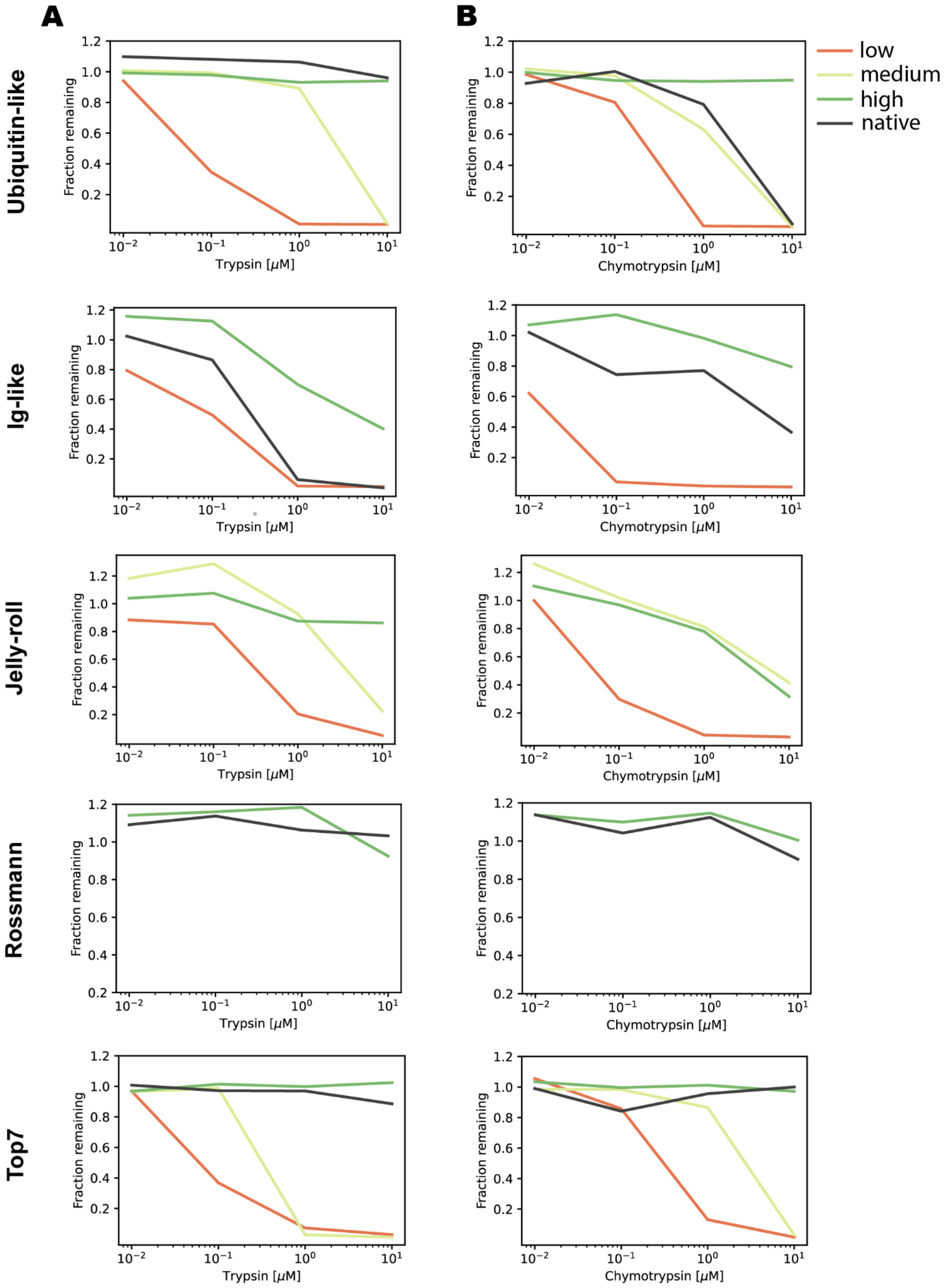


**Sup. Fig. 10 | Comparison between designs with low, medium and high protease resistance to a native protein with the same fold.**

Comparison of protease resistance on yeast for a hyperstable, stable and unstable design, and a native protein with the same fold. The designs with different levels of protease resistance were selected from the high-throughput experiment (Fig. 2). For the Jellyroll, the native protein did not express, and for the Rossmann fold the high and low resistance and unstable designs did not express. **A:** Using Trypsin protease. **B:** Using the Chymotrypsin protease.

#### Sup. Figure 11.

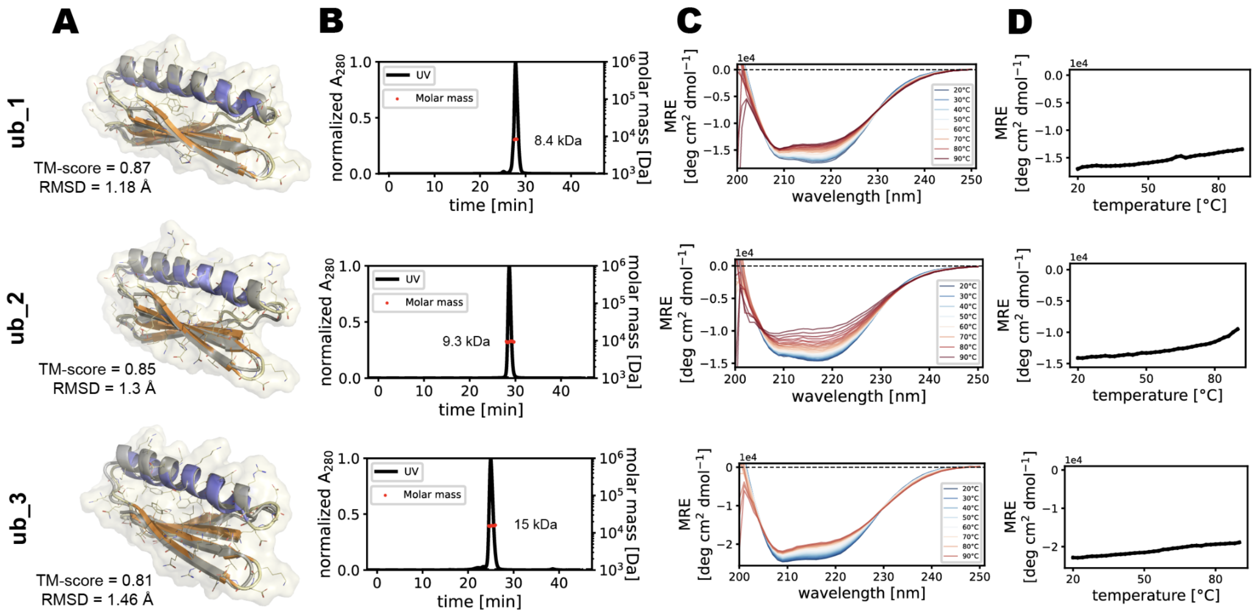


**Sup. Fig. 11 | Experimental characterization of ubiquitin-like fold designs.**

**A:** Structural model of the design for the ubiquitin-like fold (color: Genesis model; gray: AF model). **B:** SEC-MALS elution profiles showing the oligomeric state of the designs in solution. Red dots show the molecular weight as determined by SEC-MALS. The theoretical molecular weights for a monomer of the designs are: ub_1: ~7.5 kDa, ub_2: ~7.5 kDa, ub_3: ~7.3 kDa. **C:** Circular dichroism spectroscopy at different temperatures showing that the designs adopt folded structures in solution. **D:** Thermal denaturation circular dichroism spectroscopy with melting temperatures ($T_m$).

#### Sup. Figure 12.

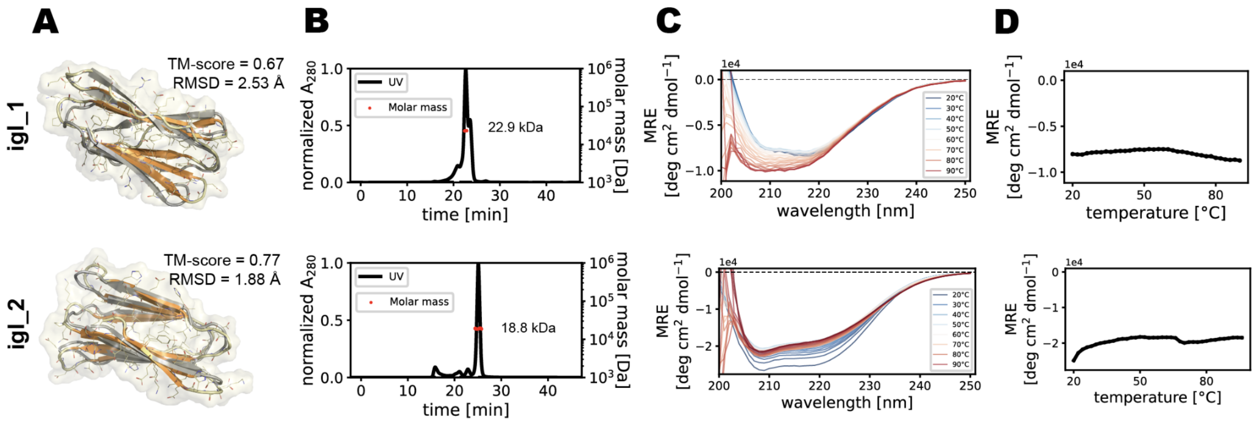


**Sup. Fig. 12 | Experimental characterization of Ig-like fold designs.**

**A:** Structural models of the designs for the Ig-like fold (color: Genesis model; gray: AF model). **B:** SEC-MALS elution profiles showing the oligomeric state of the designs in solution. Red dots show the molecular weight as determined by SEC-MALS. The theoretical molecular weights for a monomer of the designs are: igl_1: ~8.4 kDa, igl_2: ~8.5 kDa. **C:** Circular dichroism spectroscopy at different temperatures showing that the designs adopt folded structures in solution. **D:** Thermal denaturation circular dichroism spectroscopy with melting temperatures ($T_m$).

#### Sup. Figure 13.

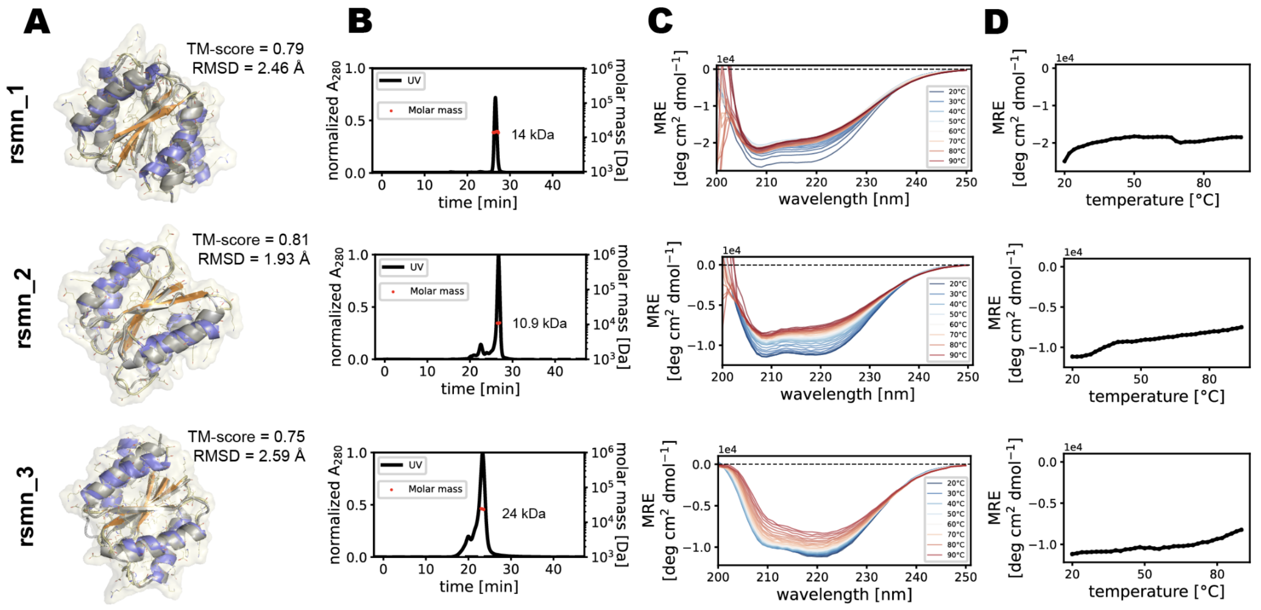


**Sup. Fig. 13 | Experimental characterization of Rossmann-like fold designs.**

**A:** Structural models of the top designs for the Rossman-like fold (color: Genesis model; gray: AF model). **B:** SEC-MALS elution profiles showing the oligomeric state of the designs in solution. Red dots show the molecular weight as determined by SEC-MALS. The theoretical molecular weights for a monomer of the designs are: rsmn_1: ~12.9 kDa, rsmn_2: ~12.5 kDa, rsmn_3: ~13.2 kDa. **C:** Circular dichroism spectroscopy at different temperatures showing that the designs adopt folded structures in solution. **D:** Thermal denaturation circular dichroism spectroscopy with melting temperatures ($T_m$).

**Sup. Figure 14.**

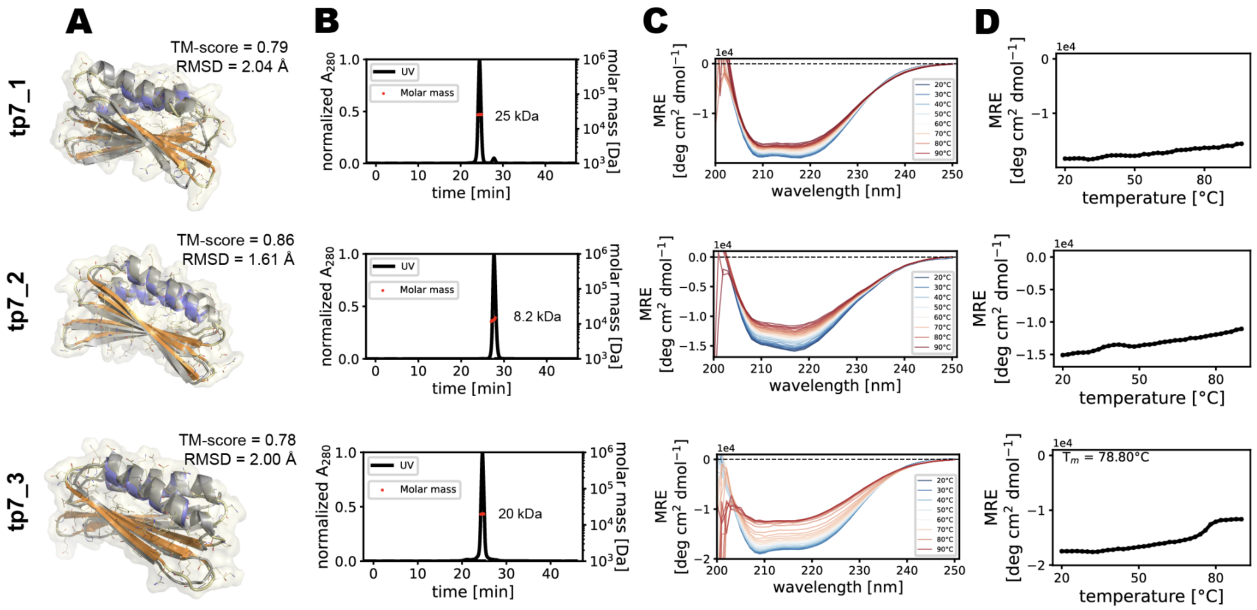


**Sup. Fig. 14 | Experimental characterization of Top7-like fold designs.**

**A:** Structural models of the top designs for the Top7-like fold (color: Genesis model; gray: AF model). **B:** SEC-MALS elution profiles showing the oligomeric state of the designs in solution. Red dots show the molecular weight as determined by SEC-MALS. The theoretical molecular weights for a monomer of the designs are: tp7_1: ~11.4 kDa, tp7_2: ~11.2 kDa, tp7_3: ~11 kDa. **C:** Circular dichroism spectroscopy at different temperatures showing that the designs adopt folded structures in solution. **D:** Thermal denaturation circular dichroism spectroscopy with melting temperatures ($T_m$).

**Sup. Figure 15.**

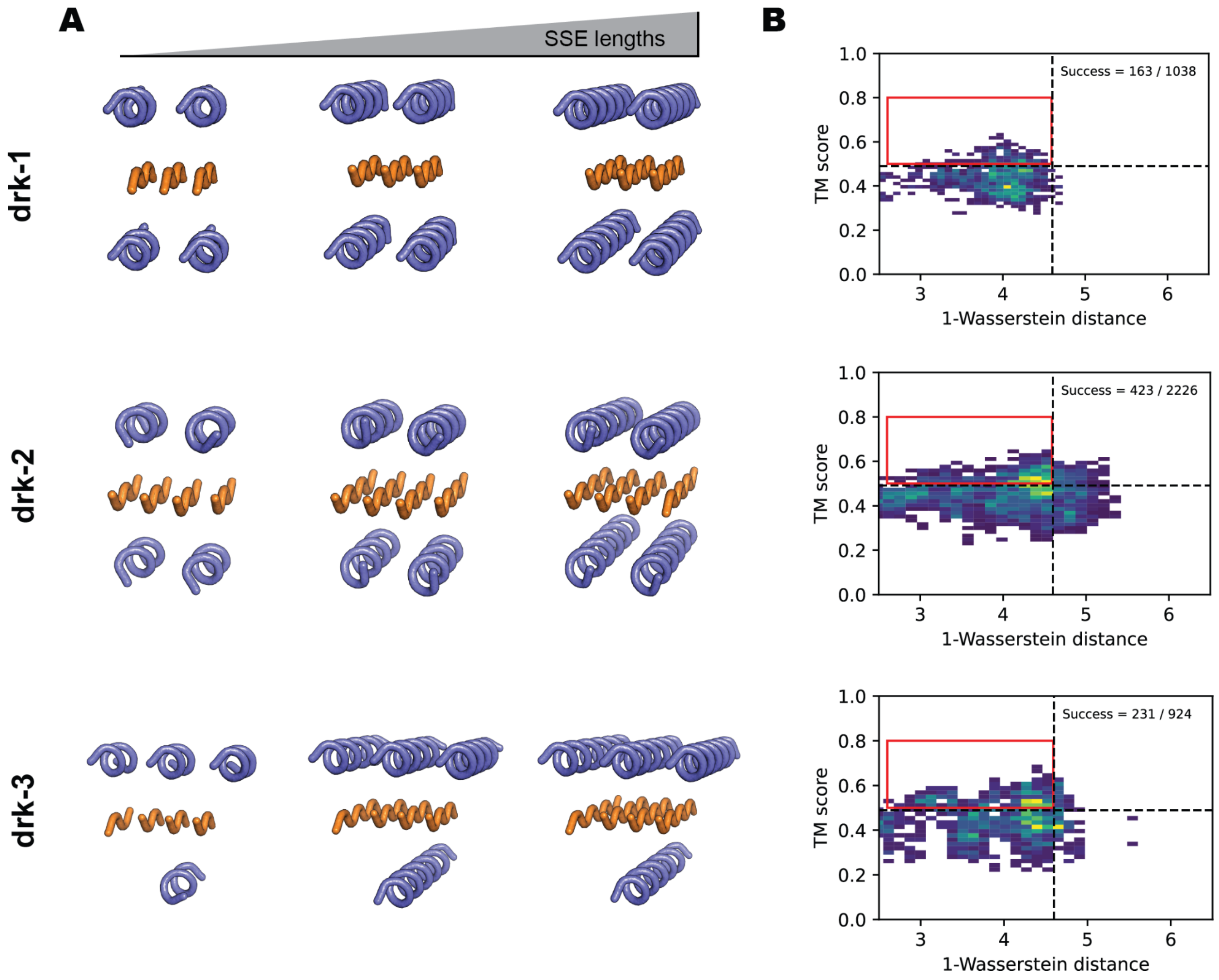


**Sup. Fig. 15 | SSE size configurations for darkfolds span a wide range of lengths**

**A:** Examples of different SSE length combinations sampled during the exploration stage to find designable candidates. As shown, a wide range from small to large SSE were searched for potential designability. **B:** Accessing backbone designability of candidates. First, the TM-score between the Genesis model and the input Sketch (> 0.5) indicates if the generated model retains the correct fold and the averaged 1st Wasserstein distance between the Genesis and trRosetta feature maps, and Genesis model feature maps indicating consistency between denoised features through the design stages.

**Sup. Figure 16.**

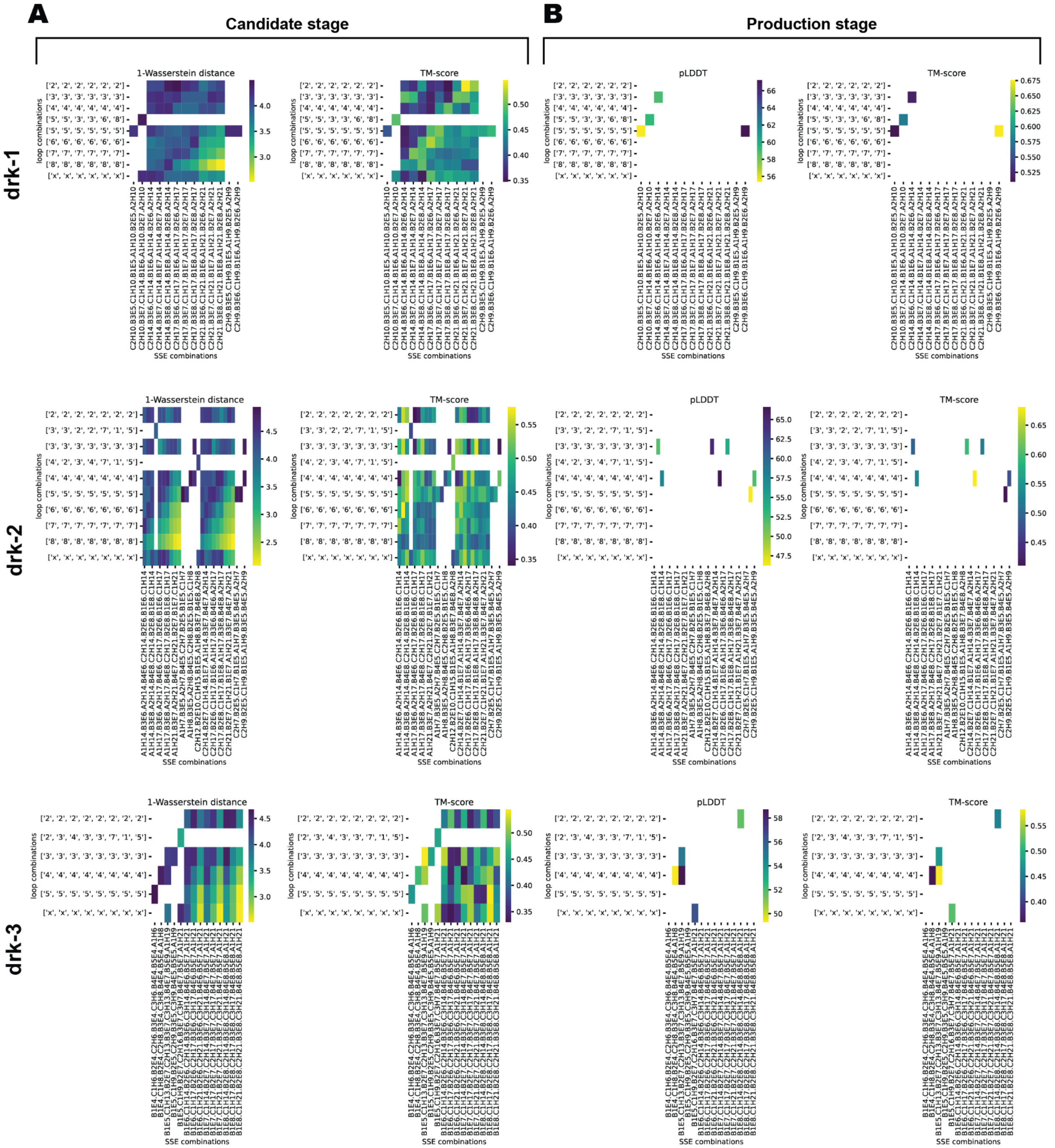


**Sup. Fig. 16 | Selective scores for combinations for candidate- and production stage for darkfolds.**

**A:** Computational scores (1-Wasserstein distance and TM-score metric) of designs for different loop and SSE combinations sampled at the candidate stage. Candidates with a 1-Wasserstein distance below 4.6 and a TM-score above 0.5 were considered for the production stage. **B:** Computational scores (pLDDT and TM-score) of upsampled designs from the candidate stage used as selection criteria.

**Sup. Figure 17.**

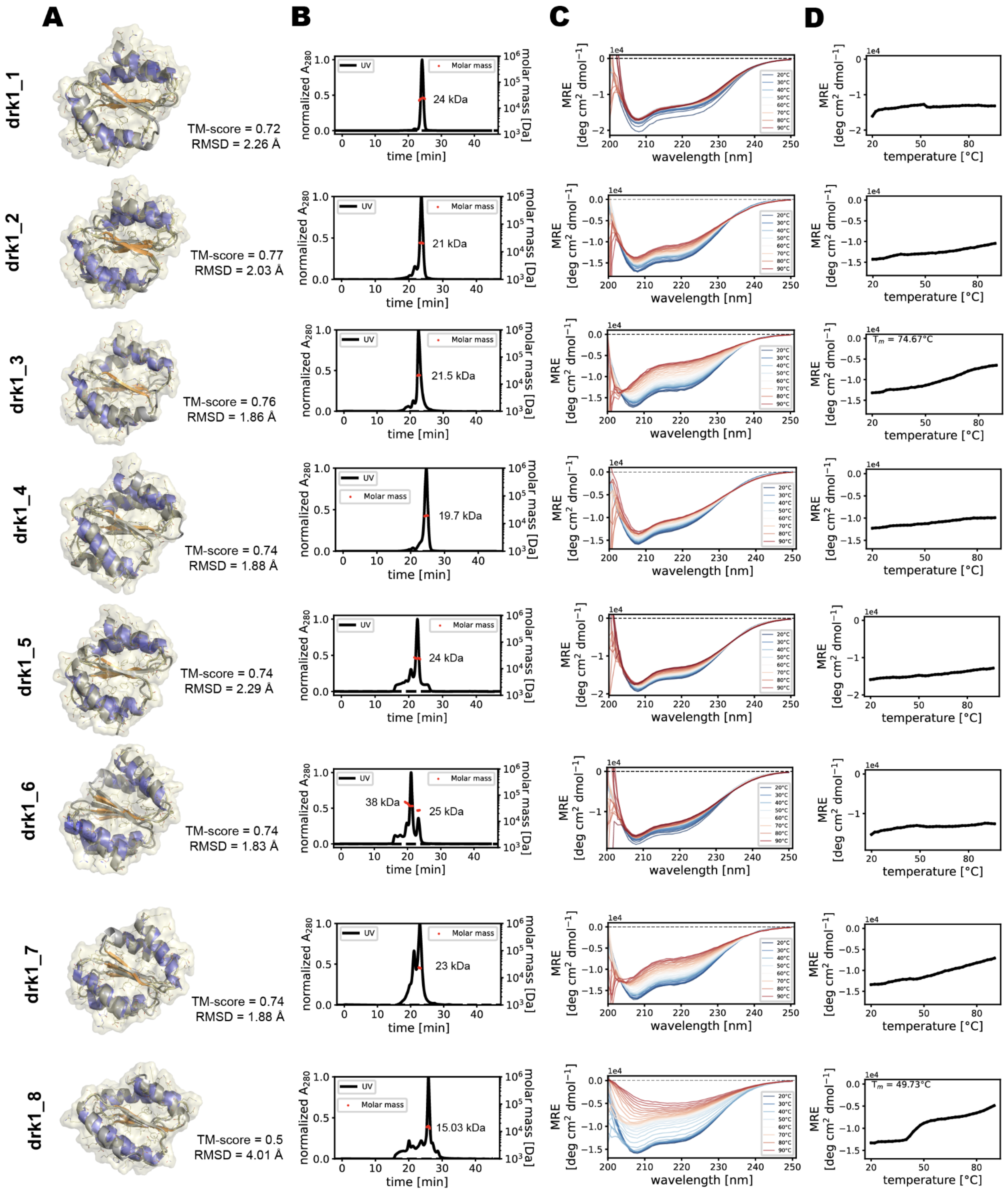


**Sup. Fig. 17 | Experimental characterization of drk1 designs.**

**A:** Structural models of the top designs (color: Genesis model; gray: AF model). **B:** SEC-MALS elution profiles showing the oligomeric state of the designs in solution. Red dots show the molecular weight as determined by SEC-MALS. The theoretical molecular weights for a monomer of the designs are: drk1_1: ~11 kDa, drk1_2: ~10.8 kDa, drk1_3: ~10.7 kDa, drk1_4: ~11 kDa, drk1_5: ~10.7 kDa, drk1_6: ~11 kDa, drk1_7: ~10.7 kDa, drk1_8: ~11 kDa. **C:** Circular dichroism spectroscopy at different temperatures showing that the designs adopt folded structures in solution. **D:** Thermal denaturation circular dichroism spectroscopy with melting temperatures ($T_m$).

#### Sup. Figure 18. - part 1

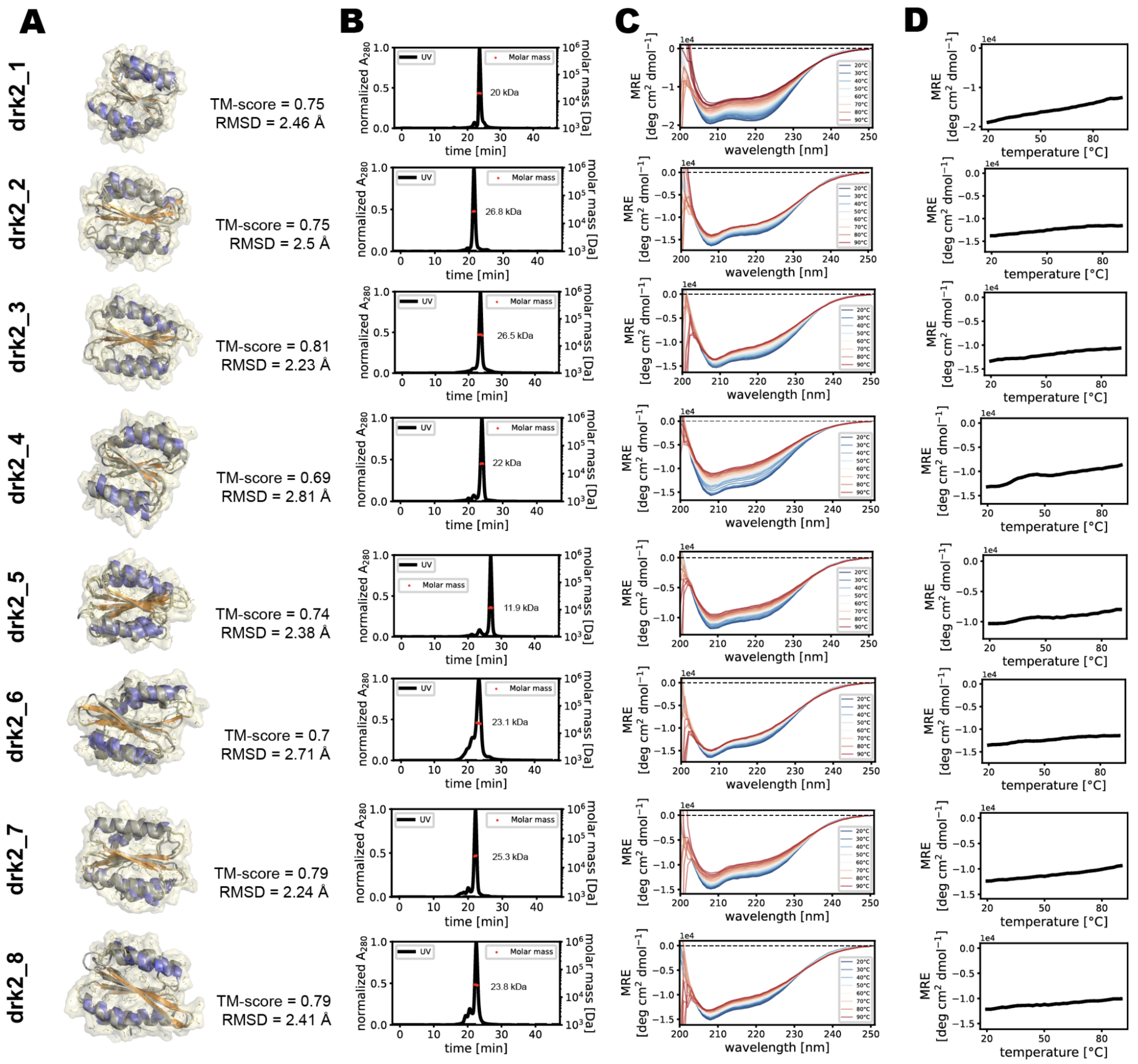

#### Sup. Figure 18. - part 2

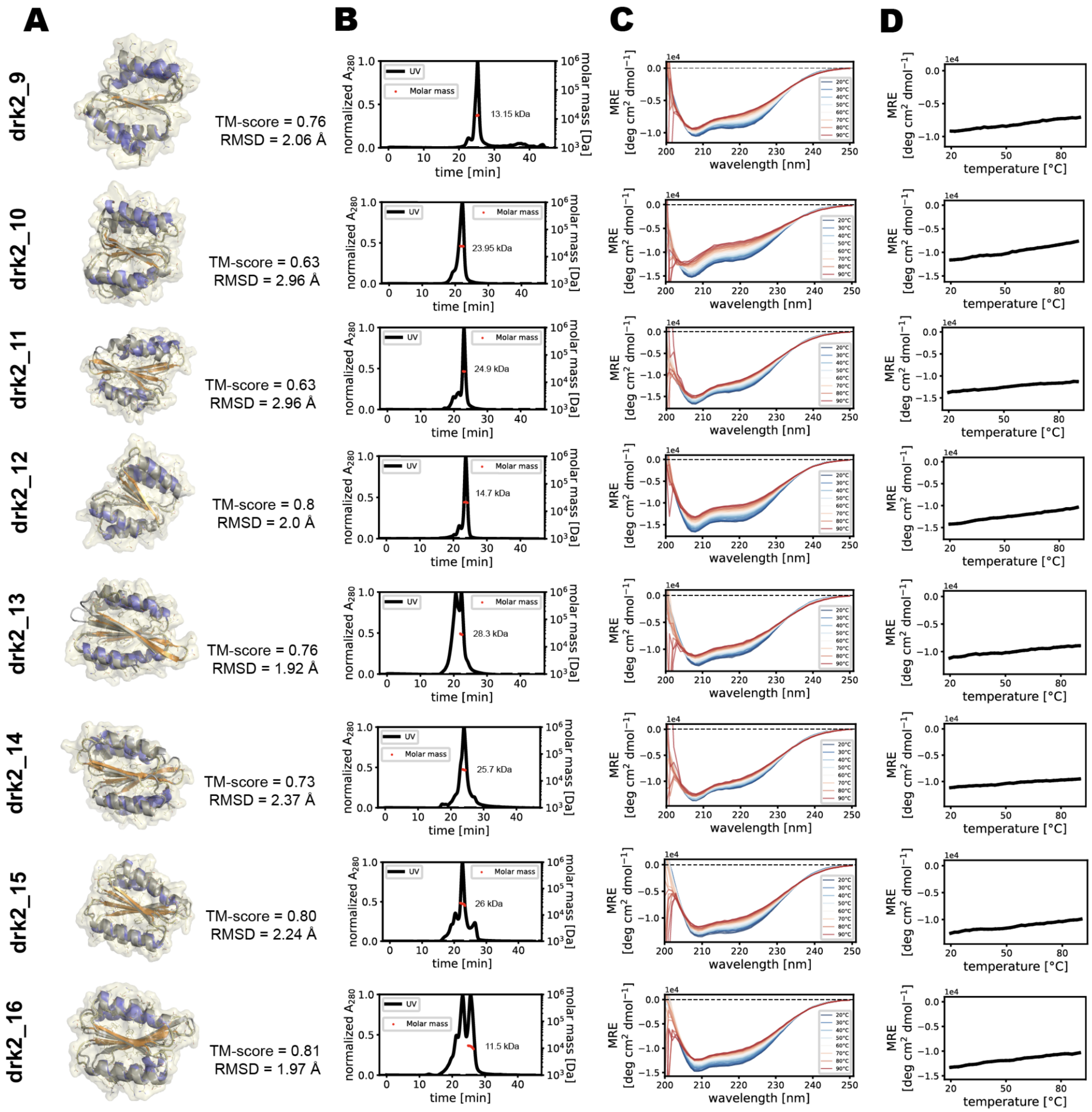


**Sup. Fig. 18 | Experimental characterization of drk2 designs.**

**A:** Structural models of the top designs (color: Genesis model; gray: AF model). **B:** SEC-MALS elution profiles showing the oligomeric state of the designs in solution. Red dots show the molecular weight as determined by SEC-MALS. The theoretical molecular weights for a monomer of the designs are: drk2_1: ~11.9 kDa, drk2_2: ~13.6 kDa, drk2_3: ~14 kDa, drk2_4: ~12 kDa, drk2_5: ~13.7 kDa, drk2_6: ~13 kDa, drk2_7: ~13.5 kDa, drk2_8: ~13.8 kDa, drk2_9: ~10.9 kDa, drk2_10: ~11.7 kDa, drk2_11: ~12.4 kDa, drk2_12: ~11.4 kDa, drk2_13: ~13.8 kDa, drk2_14: ~12.5 kDa, drk2_15: ~14.1 kDa, drk2_16: ~14.2 kDa. **C:** Circular dichroism spectroscopy at two different temperatures showing that the designs adopt folded structures in solution. **D:** Thermal denaturation circular dichroism spectroscopy with melting temperatures ($T_m$).

**Sup. Figure 19.**

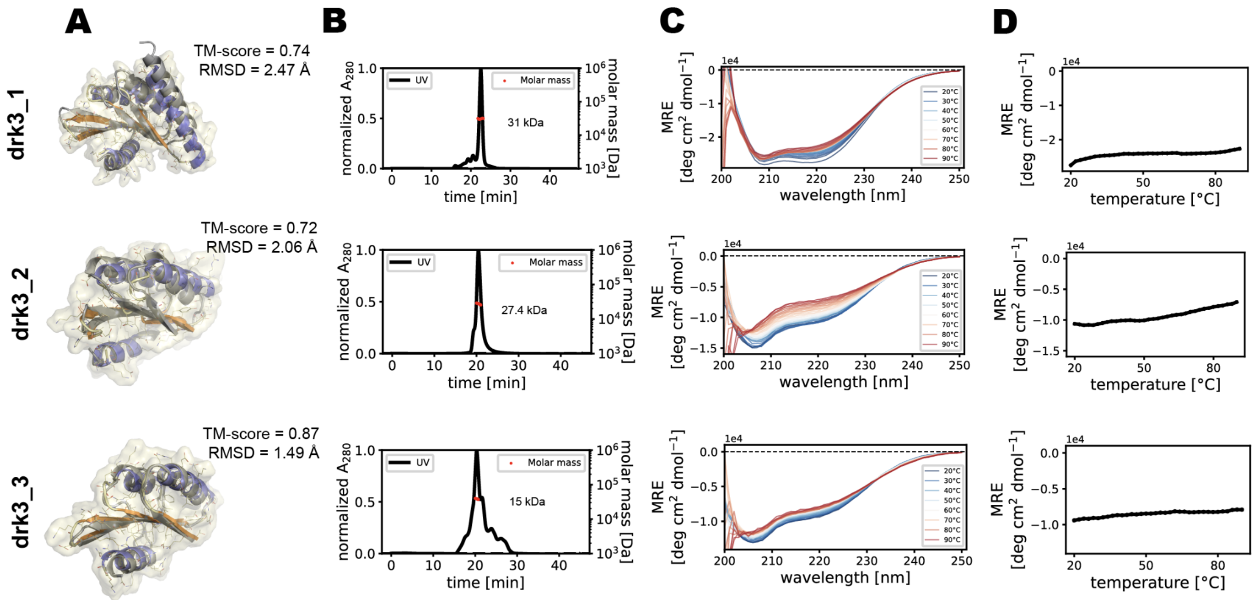


**Sup. Fig. 19 | Experimental characterization of drk3 designs.**

**A:** Structural model of the top design for each native fold (color: Genesis model; gray: AF model). **B:** SEC-MALS elution profiles showing the oligomeric state of the designs in solution. Red dots show the molecular weight as determined by SEC-MALS. The theoretical molecular weights for a monomer of the designs are: drk3_1: ~13.5 kDa, drk3_2: ~9.6 kDa, drk3_3: ~9.4 kDa. **C:** Circular dichroism spectroscopy at different temperatures showing that the designs adopt folded structures in solution. **D:** Thermal denaturation circular dichroism spectroscopy with melting temperatures ($T_m$).

### Supplementary Tables

#### Sup. Table 1.

| id | name | sequence | fold |
|---|---|---|---|
| ub_1 | N366 | VITIHVTSKDGTKTIQVSDESDDRAKEIAQKIIEELKREFKKKGIDIKIKVQYTTKDSTEEREYS | ubi |
| ub_2 | N354 | EVTIHVTSEEGTETYHVSVESDDKAEELAKKVVKEIKEEWEKRGIEVTITVHITKKDTTETKTYK | ubi |
| ub_3 | N375 | ETEIEVTKKDGTEKTKVSDESDDKAKKIAHEIIKEIKKEAKKQGIDITITIHTTKKDSTETETYS | ubi |
| tp7_2 | N418 | FRVEITVQKRDGESETRRQEFGNDSDKAEEWVKKVAKEFWDDGELSIEVKVEVDGNGNEILKKVVKALKQVVDDSGHEIKVRQETDHSGKTRVTVKVQ | top7 |
| tp7_1 | N403 | WKITITVEKRNGEKETQEYNVGDDADYARKVVKEVIERFLRDGDVSIEVTVTLDGNKDKIIHEVVKAFKEVVDDRGHEVKREQRQHEDGSIQVTVRLK | top7 |
| tp7_3 | N412 | LQVTITVQPKDGKRETRTYTVGDDTDRAKEYVEKALREFIKDGPLSVEVEVTVDGNKDKALQKVVEAAHAVVDHLLGSTEVQQHQHEDGSIQVKVKLI | top7 |
| rsmn_1 | N515 | DLLIVVSTGDTEIWKKYTEEVRRREGGPLELILYQVDPRDKEKAKKVVQEAIKQNSGTIIFFIVVGEESEELAKAIKEAVERVVDIPVIILTFSDVKDAIKQAYKLIKEYRK | rossmann |
| rsmn_2 | N5 | ILFIFYSDETEKLFEEIKKKSQKEKQFEFHKFEFTSSEEAKKIIKEIIKRWGEEEIIVIVITHDPRQEEAAKEAAKEVPAQKVIVVKVDGNDEKWKEKIKELIEK | rossmann |
| rsmn_3 | N20 | FFIIVISDGTTKWLHELIEIWRRRYKGPIELITDKIDPRDEKKIRELAHEFAKRVSGKIVFLIYIGDETHRIAEVVEEALRKVLPVPVILLKFSDPKDAIEIALKLIEKYLK | rossmann |
| jelly_1 | N314 | QTYTIQVGNKTESLQLTITLRPRTSVTLRVQGKDKSVEIHVQIHAGDQTKEWTYHLKRGNDTVEIELDHEGEITIRIRVKDTTATLTVQEI | jelly |
| igl_2 | N456 | ISWEFHEGTNSYEVTLRTDGGWQIEIHVQTSGDESHTWTLTTDTSITVRYSSDEARITLTLQDEDHEKTYTVW | iglike |
| igl_1 | N17 | FEIEVQQHGDTYEVRLKTEPNATIRITITSENGQTFTENKEPTEKITVHVSSGKVEIELRITTKDGTYTNKYK | iglike |
| drk2_5 | D26 | IKLQEVIEEYVRKYKDEQLIFFFLITRDETAEKYAQEARKTAHKLGVEEVRIIKLNDDRSIEEILKKIEEVARKVPHGKVVIFILKLHENSSYKLVVITITSDREKQLEEALKKLE | dark_2 |
| drk2_9 | D106 | KEELKRILEEVYRKTETTVIFFFPDTIAKEVKEIAHKLLHPKNNIYIFTSPTDQKAIKRVAEAIKRFPYWVLIIYDSDFKKAVEDAEELAKK | dark_2 |
| drk3_1 | D82_2 | NEELKKQTDKLWEKLKEILKKTKTESVTIEVSGDHAAEIVKLALEYLKNGLVTVHVTDGNAHELIKEIKKEAKKGLIIFVHSRSGDETKLEVYIVATSEEEEERLRRWAEEIARRLK | dark_3 |
| drk3_3 | D563 | TQYEISCGGESEERIRKLAEQSKSGEKIVLILRCGGDDKIVKIARELFSGPICIVTEKKDSCEVHFDGDESEVEKRCREKCQK | dark_3 |
| drk3_2 | D561 | EKYSVSKNEDTAREIAERAKRKKDTKVTIEFNGDDDAYRAIEEIVKKEAGAAIVLVIKGNNHLKQVRVFGEDVKEIAKELQKKI | dark_3 |
| drk2_1 | D3 | IEELAHIVLELAKQGIKVVILIFYPTVYQKLQKILKELKIEALLQIIVVPKTDKEAIREWIERLAENAQLILFLTEGRVIQIENTNTKARQEYEELIRKLQ | dark_2 |
| drk2_15 | D8 | MVKAEEVVEEVWHRYKHHKVLFILFVTHTEDAKKWAKIAKKRLHELGVEEVRIIELEDEESWKKAIEYVQKQIKKTKDGYIVIFFIIKQENSSFKIFILVLTTDHEKQLKELEEKLE | dark_2 |
| drk2_3 | D21 | MYEVKEIAEEVARRYKDTKLTFIFFVTNDETAKRIAKEAAKLLHKLGVERVEIYELNTEESLKKILKKFKEWLQKSEDGLWIVFFIIIHENSSYSIWWLISGTDEKELAEKIYKHLK | dark_2 |
| drk2_4 | D24 | MIEEVKKYIKEALKKGQPLLIIIFHDSKIQKEVEEALREVEGDGKKYKTFTANDKENVQEIWRRVARDPGWLLFIFENEVYIFKVTGSDAEKYLHELAKKYA | dark_2 |
| drk2_6 | D27 | MVEEVLKKLEERLRKQKDIVVIVLVGRTAKEKVKEVLQRVKEKVRIEVYELITTSEQLEELVKIAQKLLGGEVWIFFQVGNDSFYVIEIEKDRKEEAERQAKKYIKGSW | dark_2 |
| drk2_16 | D29 | MKKFEEYIEELARKYKETDILFIILVSKTEDLRELAQQAARIAHEIGIKEVIIIEIKTEENLQRATKIAEEIIKKTNSGIIFLFIISKTDNSSFKVYYLTLPSDREKEIEEYIKKARGSW | dark_2 |
| drk2_8 | DN21_2 | YEVKEIAEEKARRYKDTKLTFIFFVTNDETAKRIAKEAAKLLHKLGVERVEEYELNTEESLKKILKKFKEQLQKSEDGLWIVFFIIIHENSSYSIWRLISGTDEKENAEKIYKNLK | dark_2 |
| drk2_13 | DN21_3 | YEVKEIAEEKARRYKDTKLTFIFFVTNDETAKRIAKEAAKLLHKLGVERVEGYELNTEESLKKILKKFKEQLQKSEDGLWIVFFIIIHENSSYSIWRLQSGTDEKENAEKIYKNLK | dark_2 |
| drk2_10 | DN24_7 | IEEVKKGIKEALKKGQPLLIIIFHDSKIQKEVEEALREVEGDGKKNKTQTANDKENVQEIWRRVARDPGDLLFIFENEVYKEKVTGSDAEKYLHERAKKYA | dark_2 |
| drk2_11 | DN27_4 | VEEQLKKLEERLRKQKDIVVIVLVGRTAKEKVKEVLQRVKEKVRIEVDEEITTSEQLEELVKIAQKLLGGEVWIFFQVGNDSFYVIEIPKDRKEEAERQAKKKIK | dark_2 |
| drk2_12 | D199 | AEEAARIAAEWLRRGKEVVLIVLHDTEAERIEKILRKLKVKALSDILKVKQTDKETIKRAIEEALKKFDEIVVVQKGSIITISNKGNDAKKKAEELARQLE | dark_2 |
| drk2_14 | DN27_9 | VEERLKKLEERLRKQKDIVVIVLVGRTAKEKVKEVLQRVKEKVRIEVYEKITTSEQLEELVKKAQKLLGGEVWIFFQVGNDSFKVIEIRKDRKEEAERQAKKRIK | dark_2 |
| drk2_7 | DN29_1 | KKFEEYIEELARKNKETDILFIILVSKTEDLRELAQQAARIAHEIGIKEVIINEIKTEENLQRATKKAEEQIKKTNSGIIFLFIISKTDNSSFKGPELTLPSDREKEIEEYIKKAR | dark_2 |
| drk2_2 | DN29_3 | KKFEEDIEELARKYKETDILFIILVSKTEDLRELAQQANRIAEEIGIKEVEIIEIKTEENLQRATKIAEEKIKKTNSGIIFLFIISKTDNSSFKVEDLTLPSDREKEIEEYIKKAR | dark_2 |
| drk1_8 | D14 | DEELAKRAEKLAKDGDLIIFVFFDEEDARKIAEKIKKYVEQTLGSVYVIVGPTEEALKIAKEIFKKHNFKLIFLWFTSDPRQETKIKQAKKHIQ | dark_1 |
| drk1_2 | D19 | KEEIERVLREIAEKQDVTLVVFLSDTLVEEAKEAAKRVWHPKHNIVFVTGPDDERVVKEFVKAWKKYPGWVWIFIDPSAKELKEKIEEAVKK | dark_1 |
| drk1_1 | D103_1 | KEEIKRIIEEIYKKEKVTIVFLVASTLAEEVKEIAHRVLQPDQQIYIFTDPNDERVWREIAEAFKKYPSIILIFLDSDAQELAKKVEEWAKK | dark_1 |

| | | | |
|---|---|---|---|
| drk1_5 | D126_1 | KEEIEKIIKELVEKKDTNLIFFVADSLAKEVEEVAKEALQPKQNIWIITSPNDEEAIKQFVEIVKKIPVDFLIFVDDDLKELAKKIEELLKK | dark_1 |
| drk1_6 | D108_1 | KEEIERYLREIIERSNTSIILFLPDTLAKEAEEVLKKILREDQQIYLFSSNNDEKAVHEAAEAVKKIPAYLIIFYDSSFEELVKKLEKWIKQ | dark_1 |
| drk1_4 | D139 | KEEIERIFREILEKKNVTVFVILSPTKAEEAKEVIKRLIREDSEVYVFTGPDDERYAKHVVEALKKIPIEFWIIIDPDNRELAKKIKEAVHK | dark_1 |
| drk1_3 | DN19_4 | KEEIERELREIAEKQDVTLVVFLSDTLVEEAKEAAKRVWHPKHNIKDVTGPDDERVVKEFVKAWKKYPGGVRIFIDPSAKELKEKIEEAVKK | dark_1 |
| drk1_7 | DN19_7 | KEEIERELREIAEKQDVTLVVFLSDTLVEEAKEAAKRVWHPKHNIVQGTGPDDERVVKEFVKAWKKYPGWVKIFIDPSAKELKEKIEEAVKK | dark_1 |

**Sup. Tab. 1 | Experimentally characterized sequences.**

### Sup. Table 2.

| name | fold | stability | sequence |
|---|---|---|---|
| 1ubq | Ubiquitin-like fold | native & hyperstable | QIFVKTLTGKTITLEVEPSDTIENVKAKIQDKEGIPPDQQRLIFAGKQLEDGRTLSDYNIQKESTLHLVLRLRGG |
| n362 | Ubiquitin-like fold | hyperstable | EIQIHVTTKTGTETYKVSDDSDHKAKEKAQKIIKELQEKYKKQGIDVTITVELTKKDSTEKETYS |
| n371 | Ubiquitin-like fold | stable | QIQLHVTTKEGTETYTISDESDDKQEKLAQELIEKIRREFQKKGIEVTITLETTKKDSTEKKTYQ |
| n377 | Ubiquitin-like fold | unstable | ISVEVRVTNDSSETQSYSFSQERYDGQDELLERWAQELVKRIQKQLQPKGSSWRQETQQEGNEKKIELEI |
| 1cyl | Ig-like | native & hyperstable | PSVFIFPPSDEQLKSGTASVVCLLNNFYPREAKVQWKVDNALQSGNSQESVTEQDSKDSTYSLSSTLTLSKADYEKH KVYACEVTHQGLSSPVTKSFN |
| n466 | Ig-like | hyperstable | LSFSINEGTDSVQLTVTTEGGWQVTLEIQTRGDEKQQYTITGDTSLTIHLSKHEAKLTVTLQDETHEETHTLI |
| n474 | Ig-like | stable | FEVRLNSGTKSITITVKVEGGWQIEIKVETSGDQTYKVTLTGDTEVQVTVSEQPARFTVTVQDEDHEKTITLL |
| n486 | Ig-like | unstable | YELRKQQHGSTYTVEVRSEKDAKIHIEIQAEKGQKFTENVQPEGRSRITVTEGSAEVRLEIHKKDGTRSNTLE |
| 4nia | Jellyroll | native | TWVRAIPFEVSVQSGIAFKVPVGSLFSANFRTDSFTSVTVMSVRAWTQLTPPVNEYSFVRLKPLFKTGDSTEEFEGRA SNINTRASVGYRIPTNLRQNTVAADNVCEVRSNCRQVALVISCCFN |
| n344 | Jellyroll | hyperstable | TLEYTIEGSSLQIHVPGGGGSLTLEVTKDAKLTLEKGRNSETKEAKDSLKIQVSLSTDITVRVEISGNLTFKVE |
| n329 | Jellyroll | stable | QKVTVELGNSTSSITITVTLDKHDRIQIEVELDDDQVEIEVHVHAGDETKTWHLRGTKGNDKVSIELLENGEVTLTIRWL KSHGRITLQNL |
| n346 | Jellyroll | unstable | KIKSKIPGSSVTFRVPGGGGTITVRVQKDATVTIENGRSSQTYTGDKHIEVTVTINDDIRIHVEIKGNLEVEIK |
| 1pdo | Rossmann | native & hyperstable | TIAIVIGTHGWAAEQLLKTAEMLLGEQENVGWIDFVPGENAETLIEKYNAQLAKLDTTKGVLFLVDTWGGSPFNAASRI VVDKEHYEVIAGVNIPMLVETLMARDDDPSFDELVALAVETGREG |
| n501 | Rossmann | hyperstable | ILIIIISTDDTEEWKKATKEWQKEQGGPIKLVLRQVDPRDKEKAKELAKEIAKKNSGEIIFIVVLGDENKQLAEEIHKALRKL IPLPVYVIKVSDLKDAIKQVWEIIQKIYR |
| n522 | Rossmann | stable | TVLFVLSEDDKKEWEEWVKRTKKEEEWEIVTLEFKETDQAEEVWKKAIKKYGDDTLYLIVFFSDPRLQEAAERAVKEA KLDKWILVQTKNNEEEAKEIVKREIKK |
| n530 | Rossmann | unstable | IVVIVVFSTDDEDKIKKWKEIVEKLLKGGGKTFEIWFLVVNGSEAEEILKRYIKKAHQYGKLVLIVQGDSEIQKAAKKAHE LAKKVEVILLLISIDGDESKAEKLAKEAWERIK |
| 1qys | Top7 | native & hyperstable | GDIQVQVNIDDNGKNFDYTYTVTTESELQKVLNELMDYIKKQGAKRVRISITARTKKEAEKFAAILIKVFAELGYNDINVT FDGDTVTVEGQLEGGSLE |
| n418 | Top7 | hyperstable | FRVEITVQKRDGESETRRQEFGNDSDKAEEWVKKVAKEFWDDGELSIEVKVEVDGNGNEILKKVVKALKQVVDDSGH EIKVRQETDHSGKTRVTVKVQ |
| n435 | Top7 | stable | NFTLKTEDYEQEFKVTSGTKDKLKKVLEEIIRKAHQFPKPVEIEIEVEKEISEELAQEFKKVAEKLLPNYKVELQTESKDR ITLTVH |
| n429 | Top7 | unstable | NVEFTFENNQQKFSLTSGKGDEALKIIEEIIRKLQSVPTPIEIKLTWEPEISEEFARRAKEIAKRLWPNYEIRDQQENDNR QTFHLL |

**Sup. Tab. 2 | Sequences from protease resistance comparison to native proteins with the same fold.**
